# The RNA binding proteins Ddx6 and Ddx61 support development in mRNA-decay deficient *pnrc2* mutants

**DOI:** 10.1101/2025.05.08.652725

**Authors:** Thomas L. Gallagher, Monica C. Blatnik, Clare C. Austin, Kathryn G. Thompson, Danielle M. Pvirre, Ryan Denniston, Angelina Morgan, Michael G. Kearse, Sharon L. Amacher

**Author notes:** Oberlin College of Arts and Sciences, 38 E. College St., Oberlin, OH, 44074, USA.

## Abstract

Somitogenesis is controlled by the segmentation clock, a molecular oscillator that controls periodic gene expression in unsegmented mesoderm and is regulated by a negative feedback loop driven by Hes/Her transcriptional repressors. In zebrafish, Pnrc2 is required for decay of *her1* transcript and additional oscillatory gene transcripts. Despite accumulation of numerous mRNAs including those encoding developmental regulators, overt embryonic phenotypes are absent in *pnrc2* mutants. Our previous work suggested that accumulated mRNAs are not translated in *pnrc2* mutants, though the underlying mechanism(s) was unknown. We show here that many overexpressed transcripts in *pnrc2* mutants have shortened poly(A) tails and are disengaged from ribosomes, and that deadenylation inhibition leads to somite defects in *pnrc2* mutants. Transcripts encoding the P-body-associated protein, Ddx61, are overexpressed and engaged with ribosomes in *pnrc2* mutants, leading to an increase in Ddx61 protein that may compensate against the negative effects of transcript accumulation. Co-depletion of Ddx61 and Ddx6 in *pnrc2* mutants increases *her1* accumulation and leads to pleiotropic defects. Together, our results show that multiple post-transcriptional mechanisms ensure proper translation when mRNA decay is inhibited.

**Summary statement:** Pnrc2 promotes decay of tail-shortened, ribosome-disengaged mRNAs and co-regulates *her1* mRNA decay with Ddx6 and Ddx61, P-body-associated factors that sustain normal development in decay-deficient *pnrc2* mutants.

## Introduction

Vertebrate somitogenesis is a fundamental developmental process (Kageyama et al., 2012) characterized by the anterior to posterior sequential segmentation of the embryonic mesoderm into blocks of tissue called somites. This rhythmic process is controlled by a molecular oscillator called the segmentation clock which operates in the presomitic mesoderm (PSM) and is characterized by rapid cycles of mRNA and protein expression (Hubaud and Pourquié, 2014; Oates et al., 2012; Pourquié, 2011). The segmentation clock is regulated by a self-sustaining negative feedback loop, where a core oscillatory factor, a transcriptional repressor, inhibits expression of downstream oscillatory genes, including itself.

For proper somite formation, rates of each oscillatory step, from transcription to translation to decay, must be regulated to ensure correct somite size and number (Gomez et al., 2008; Holley et al., 2000; Keynes and Stern 1988; Lewis, 2003; Matsuda et al., 2020; Palmeirim et al., 1997; Schröter and Oates, 2010). Real-time segmentation clock reporters that recapitulate clock dynamics in vivo must not only contain critical transcriptional regulatory regions that drive oscillatory expression, but also contain features including 3′UTR sequences and protein motifs that destabilize reporter mRNAs and proteins, respectively (Aulehla et al., 2008; Delaune et al., 2012; Masamizu et al., 2006; Simsek et al., 2023; Yoshioka-Kobayashi et al., 2020).

We have focused on post-transcriptional regulation of the segmentation clock (Blatnik et al., 2023), motivated by our discovery that the zebrafish *pnrc2* gene promotes rapid decay of oscillatory gene transcripts (Dill and Amacher, 2005; Gallagher et al., 2017; Tietz et al., 2020). Transcriptional oscillations are normal in zebrafish *pnrc2* mutants, but *her1* mRNA (and other oscillatory gene transcripts) accumulate due to mRNA decay defects (Gallagher et al., 2017).

Surprisingly, despite a 4-6-fold increase in *her1* and *dlc* oscillatory gene transcripts in *pnrc2* mutants, oscillatory gene protein is expressed normally (Gallagher et al., 2017) and there is no overt segmentation phenotype. Discordance between mRNA and protein levels might be caused by inefficient translation due to binding of translational repressors and/or localization to granules like P-bodies that are sites of non-translating mRNAs (Luo et al., 2018; Ostareck et al., 2014).

To better understand the role of *pnrc2* during segmentation, we used RNA-Seq analysis and identified over 1,700 transcripts that were significantly overexpressed at least 1.2-fold in *MZpnrc2* mutants. Because *pnrc2* is expressed broadly in embryos (Gallagher et al., 2017), we anticipated that Pnrc2 likely functions in many tissues. GO analysis reveals enrichment for many processes associated with development and mRNA regulation. Motif analysis shows that overexpressed transcripts are enriched for well-known mRNA-destabilizing motifs. Several accumulated oscillatory gene transcripts have shortened poly(A) tails in *MZpnrc2* mutants, suggesting that they are normally unstable. Polysome profiling of *MZpnrc2* mutants shows that accumulated transcripts of several oscillatory genes are primarily disengaged from ribosomes.

*MZpnrc2* mutants are sensitive to deadenylation inhibition during segmentation, and later to depletion of P-body-associated factors Ddx6 and Ddx61, though whether these effects are connected remains unknown. Our results support that discordant expression of oscillatory gene mRNA and protein in *MZpnrc2* mutants is due to inefficient translation of accumulated transcripts. In the absence of *pnrc2* function, P-body-associated factors Ddx6 and Ddx61 buffer against mRNA decay dysfunction and support embryonic development, implicating a broader role for Pnrc2 and Ddx6/Ddx61 during development across many tissues.

## Results

### Pnrc2-regulated transcripts are associated with key biological processes and are enriched for mRNA destabilizing motifs

To globally identify transcripts that require *pnrc2* for proper expression, we performed RNA-Seq analysis on wild-type (WT) and maternal-zygotic *pnrc2* (*MZpnrc2*) mutant embryos at mid-segmentation stages. We detected 2,256 differentially expressed transcripts with a false discovery rate of q < 0.05 (Fig. 1A, Tables S1-S3) using Kallisto to quantify transcript abundance (Bray et al., 2016) and Sleuth for differential expression analysis (Pimentel et al., 2017). Most differentially expressed transcripts were overexpressed (n = 1,769 transcripts), as expected for an mRNA decay-deficient mutant. Among overexpressed transcripts, >99% are annotated as protein-coding by Ensembl’s biotype annotation (n = 1,759 transcripts), suggesting that Pnrc2 regulates decay of transcripts with full open reading frames rather than transcripts containing premature termination codons that are subject to nonsense-mediated decay (NMD).

**Fig. 1.**
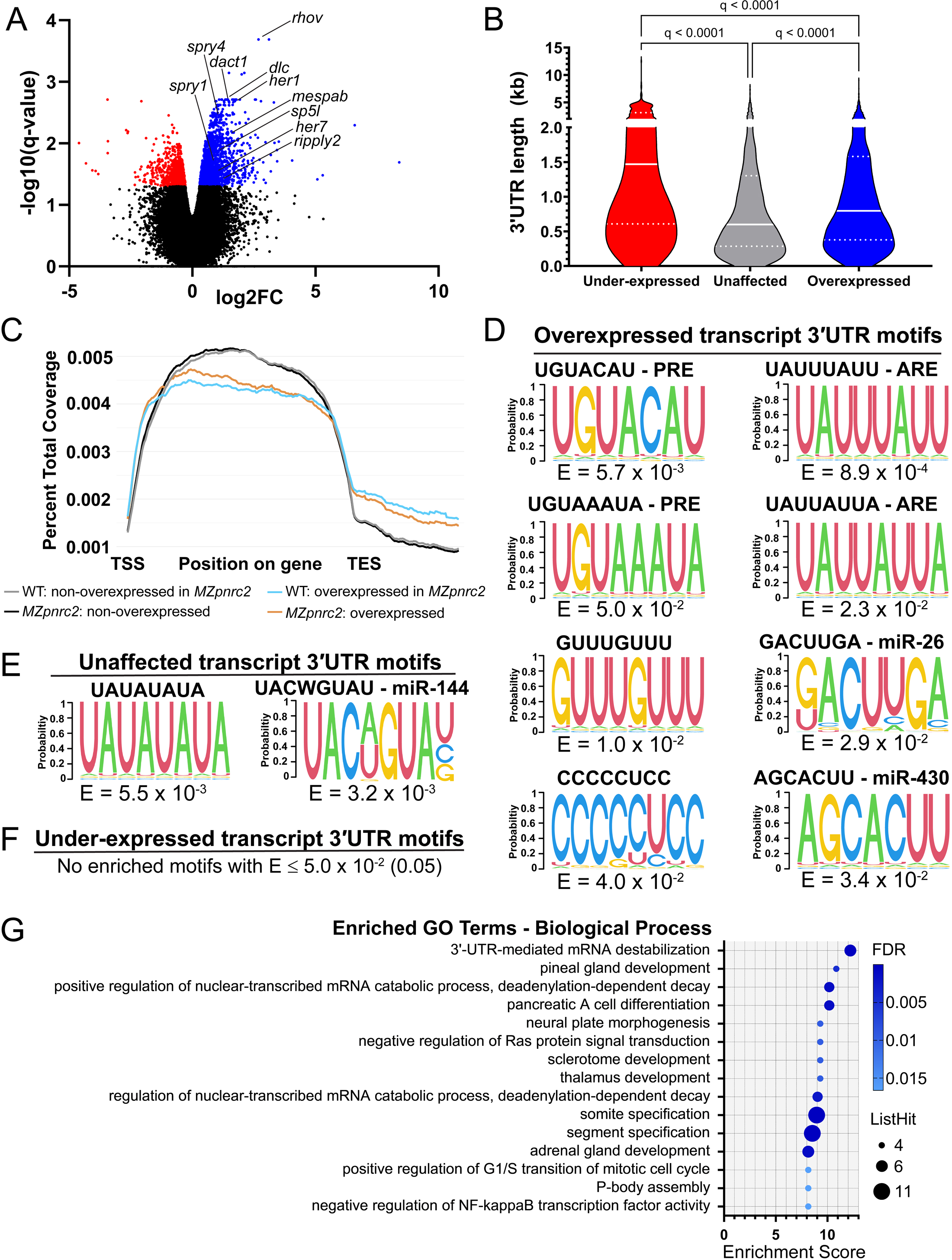
Overexpressed transcripts in *MZpnrc2* mutants have a longer 3′UTR length distribution than unaffected transcripts, are associated with many biological processes, and are enriched for destabilizing motifs. (A) Volcano plot showing transcripts that are overexpressed (q < 0.05, n = 1,769) (blue), under-expressed (i.e. transcripts with reduced expression; q < 0.05, n = 856) (red) or not significantly differentially expressed (q > 0.05, n = 32,533) (black) in *MZpnrc2* mutant versus WT embryos at mid-segmentation stages. Key segmentation-associated transcripts (Fig. 2) are indicated on the volcano plot. (B) Violin plots of 3′UTR lengths for overexpressed (q < 0.05, n = 1,333) (blue), under-expressed (q < 0.05, n = 685) (red), and unaffected (log2FC -0.05 to 0.05, n = 1,352) (grey) transcripts. (C) Metagene plot showing normalized RNA-Seq read coverage in WT (cyan) and *MZpnrc2* mutant (orange) samples for transcripts that are overexpressed in *MZpnrc2* mutants (q < 0.05) and read coverage in WT (grey) and *MZpnrc2* mutant (black) samples for transcripts that are non-overexpressed in *MZpnrc2* mutants (i.e. all transcripts that were not significantly overexpressed). (D-F) Enriched motifs in (D) overexpressed (n = 1320), (E) unaffected (n = 1334), and (F) under-expressed (n = 690) transcript 3′UTRs (E-value < 0.05) identified using STREME. (G) Significantly enriched terms within the aspect biological process using GO analysis of overexpressed genes in *MZpnrc2* mutants. Among the top 30 enriched terms (see Fig. S1C), 15 terms are shown that contain gene sets ≥5 and ListHit values ≥4. FC = fold change; kb = kilobase; TSS = transcription start site; TES = transcription end site; FDR = false discovery rate; ListHit = number of overexpressed genes per GO term.

NMD can also be induced by specific features in transcripts with full open reading frames. One such feature is the presence of an upstream open reading frame (uORF) (Mendell et al., 2004; Hurt et al., 2013; Johnstone et al., 2016). Using stringent filtering criteria (see methods), we identified uORF-containing transcripts using a list validated by ribosome profiling (Johnstone et al., 2016) and find that overexpressed transcripts in *MZpnrc2* mutants are enriched for uORFs relative to non-overexpressed transcripts (21.2% vs 15.7%, respectively, p = 2.832x10^-07^ by a Fisher’s Exact Test) (Table S3). These results indicate that Pnrc2 promotes decay of some, but not all uORF-containing transcripts. NMD can also be induced by intron- containing 3′UTRs (Schweingruber et al., 2013; Lykke-Andersen & Jensen, 2015), however, few overexpressed transcripts in *MZpnrc2* mutants contain 3’UIs (n = 50 transcripts of 1769 overexpressed transcripts) when compared to a published list of 3’UI-containing transcripts (Gangras et al., 2020). Finally, 3′UTR length can also influence mRNA stability (Schweingruber et al., 2013; Lykke-Andersen & Jensen, 2015), so we compared the length distributions of overexpressed, unaffected, and under-expressed transcript 3′UTRs (see methods). Although overexpressed transcript 3′UTRs have a significantly longer length distribution than unaffected transcript 3′UTRs, under-expressed transcripts have an even longer length distribution than overexpressed and unaffected transcript 3′UTRs (Fig. 1B), suggesting that 3′UTR length is not a primary mechanism by which Pnrc2 promotes transcript decay. To assess whether 3′UTR lengths are different between WT and *MZpnrc2* mutant embryos, sequencing reads were aligned using STAR (Dobin et al., 2013) and metagene plots were generated using deepTools (Ramírez et al., 2016). Metagene plots of gene body read coverage for WT and *MZpnrc2* mutants are very similar, including a steep drop in coverage downstream of transcription stop sites (TESs), indicating that 3′UTR length is unaffected in *MZpnrc2* mutants (Fig. 1C).

Normalized read coverage bedgraphs for the oscillatory gene transcripts *her1*, *her7*, *dlc*, and *rhov,* all of which are overexpressed in *MZpnrc2* mutants, similarly show no differences in read coverage between genotypes (Fig. S1A-B). Read coverage downstream of TESs, although very low relative to gene body coverage, is higher in both WT and *MZpnrc2* mutants for transcripts that are overexpressed relative to non-overexpressed transcripts in *MZpnrc2* mutants (Fig. 1C).

These results suggest that transcripts that depend on Pnrc2 for decay may have higher rates of Pnrc2-independent transcriptional read-through beyond their annotated TES (or alternative poly(A) cleavage site usage).

Our previous work showed that the Pnrc2-regulated *her1* transcript contains two critical destabilizing motifs in the 3′UTR, a Pumilio Response Element (PRE) and AU-rich Element (ARE) (Tietz et al., 2020). To determine whether 3′UTRs of overexpressed transcripts in *MZpnrc2* mutants are enriched for PREs, AREs, and/or additional mRNA motifs, we performed motif analysis using the motif discovery tool STREME (Bailey, 2021). Among all 8 statistically significant enriched motifs (Fig. 1D, Tables S4-S5), the sequences UGUACAU and UGUAAAUA match a PRE, a motif associated with transcript destabilization (Arvola et al., 2020; Bohn et al., 2018; Galgano et al., 2008; Gerber et al., 2006; Hafner et al., 2010; Lu and Hall, 2011; Morris et al., 2008; Murata et al., 1995; Wang et al., 2002; White et al., 2001; Zamore et al., 1997; Zamore et al., 1998). The enriched motifs UAUUUAUU and UAUUAUUA match AREs, motifs recognized by a broad group of ARE-binding proteins (ARE-BPs) that can both inhibit and promote mRNA decay and/or translation (Briata et al., 2005; Garneau et al., 2007; Gratacós and Brewer, 2010; Lykke-Andersen and Wagner, 2005; Sanduja et al., 2011). Additional enriched motifs include GUUUGUUU that matches GU-rich elements (GREs), a motif associated with mRNA decay (Rattenbacher et al., 2010), and CCC<u>CCUC</u>C that contains a destabilizing CCUC motif (Vejner et al., 2019). Lastly, we find enrichment for two miRNA target sites, the motif K<u>ACUUGA</u> that matches the miR-26 seed target site UACUUGA (Crippa et al., 2016; Icli et al., 2013; Klett et al., 2019; Watterston et al., 2019) and the motif A<u>GCACUU</u> that matches the miR- 430 target site GCACUU (Baia Amaral et al., 2024; Bazzini et al., 2012; Giraldez et al., 2006; Liu et al., 2020; Rabani et al., 2017; Vejnar et al., 2019). Overall, all significantly enriched 3′UTR motifs in overexpressed transcripts are associated with mRNA-destabilizing functions.

STREME analysis of unaffected transcript 3′UTRs reveals significant enrichment for two motifs, none of which overlap with motifs enriched in the overexpressed motif analysis (Fig. 1E, Tables S4-S5). One motif is a UA-dinucleotide repeat that in yeast is associated with enhanced poly(A) site cleavage (Graber et al., 1999). The second enriched motif, UACWGUAU, matches the miR-144 seed target sites ACAGUAU and UACAGUA (Fu et al., 2009; Su et al., 2014; Kretov et al., 2024). miR-144 is an erythroid-specific miRNA that is not expressed at mid-segmentation stages (Fu et al., 2009), which may explain why miR-144 motifs are enriched in 3′UTRs of unaffected transcripts. STREME analysis on under-expressed transcript 3′UTRs showed no motif enrichment (Fig. 1F, Tables S4-S5). Altogether, 3′UTR motif analysis shows that Pnrc2 influences decay of mRNAs enriched with destabilizing RNA motifs.

To explore possible functions of differentially expressed transcripts in *MZpnrc2* mutants, we conducted GO enrichment analysis (Thomas et al., 2022). GO analysis of overexpressed transcripts identified many significantly enriched GO terms (Figs 1G, S1C-E,Tables S4-S5) including somite specification and negative regulation of neuron differentiation, processes that rely upon *Hes/her* function for proper differentiation (Bonev et al., 2012; Hatakeyama et al., 2004; Ishibashi et al., 1995; Ohtsuka et al., 1999; Soto et al., 2020; Tan et al., 2012). Additional enriched terms are related to developmental pathways involving Fgf, Wnt, Notch, and Ephrin signaling (Fig. S1D, Table S5). Terms associated with post-transcriptional regulation are also enriched including 3′UTR-mediated destabilization, deadenylation-dependent decay, and P-body assembly (Fig. 1G, Table S5). Overexpressed genes in these categories may represent transcripts that depend on Pnrc2 for their decay or transcripts that compensate for loss of Pnrc2 function. Alternatively, overexpressed genes might be genes that are transcriptionally repressed by Pnrc2. GO analysis of under-expressed transcripts identified significant enrichment for terms including translation, peptide biosynthetic process, and eukaryotic translation elongation factor complex (Fig. S1F-H). Under-expressed genes in these categories may represent transcripts that are stabilized by Pnrc2 or genes that are downregulated as a compensatory mechanism to prevent over-translation of accumulated transcripts in *MZpnrc2* mutants. Alternatively, under-expressed genes might be genes that are transcriptionally promoted by Pnrc2.

### mRNA expression patterns of many developmental genes are disrupted in pnrc2 mutants

We characterized mRNA expression patterns and validated mRNA expression levels of select genes associated with segmentation, oscillatory expression, and/or Fgf or Wnt signaling that were identified as overexpressed in *pnrc2* mutants via RNA-Seq analysis. For in situ hybridization, colouration reactions for each probe were developed in parallel for identical reaction times in WT and *MZpnrc2* mutants. For RT-qPCR, expression levels were quantified using whole embryo extracts from mid-segmentation stage WT and *MZpnrc2* mutants. The expression pattern of *rhov*, an oscillatory gene in zebrafish PSM (Krol et al., 2011), is restricted in WT embryos but appears expanded in *MZpnrc2* mutants (Fig. 2A, A’ vs B, B’), although it is possible that the difference reflects signal detection limits rather than absence of expression.

**Fig. 2.**
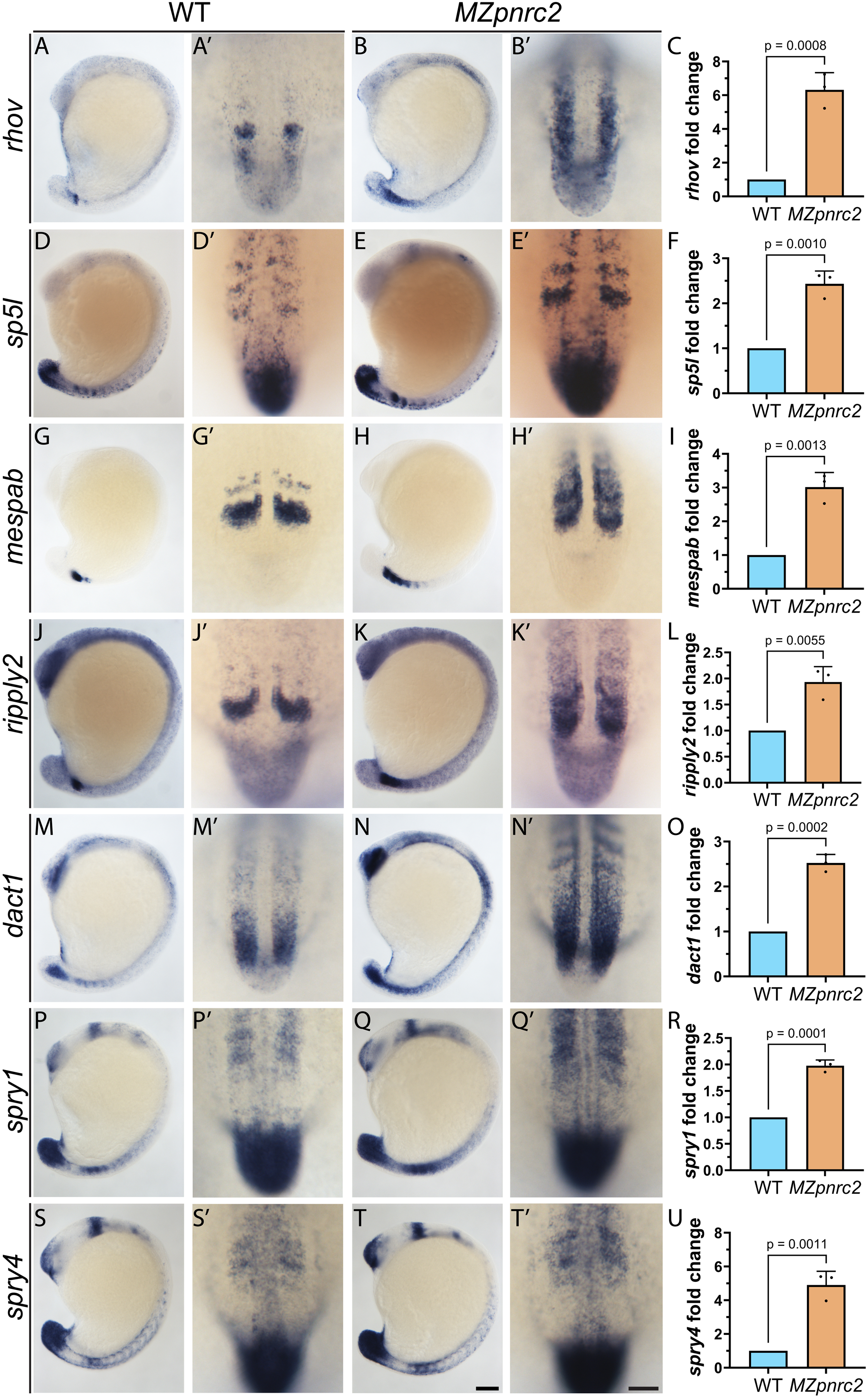
Expression of transcripts associated with segmentation and other developmental processes are disrupted in *MZpnrc2* mutants. (A-U) WT and *MZpnrc2* mutants were raised to mid-segmentation stages (14-18 hpf) and probed for *rhov* (A, A’ vs B, B’, RT-qPCR in C), *sp5l* (D, D’ vs E, E’, RT-qPCR in F), *mespab* (G, G’ vs H, H’, RT-qPCR in I), *ripply2* (J, J’ vs K, K’, RT- qPCR in L), *dact1* (M, M’ vs N, N’, RT-qPCR in O), *spry1* (P, P’ vs Q, Q’, RT-qPCR in R), and *spry4* (S, S’ vs T, T’, RT-qPCR in U). Colouration reactions for each probe were developed in parallel for identical reaction times. Representative images from each genotype are shown. In situs were performed twice independently with consistent results (n = 15-20 embryos per probe per genotype for each experiment). RT-qPCR was performed separately from in situ experiments using whole embryos (n = 15 embryos per bio-replicate per genotype). Fold change values are the mean of three bio-replicate values derived from the average of two technical replicates. P-values were calculated using an unpaired t-test. In situs are shown in lateral view (A-T), magnified in dorsal view to the right of each embryo (A’-T’). Scale bars = 100 um (A-T); dorsal view = 50 um (A’-T’). hpf = hours post fertilization.

RT-qPCR validates that *rhov* expression is significantly increased in *MZpnrc2* mutants (Fig. 2C). Expression of *sp5l* (Fig. 2D, D’ vs E, E’, RT-qPCR in F), a gene whose ortholog *Sp5* oscillates in mouse PSM (Krol et al., 2011; Matsuda et al., 2020), is similarly affected. Expression of somite polarity genes *mespab* and *ripply2* also show expanded and elevated expression in *MZpnrc2* mutants (Fig. 2G, G’ vs H, H’’ and J, J’ vs K, K’, RT-qPCR in I and L). Expression of *dact1*, a modulator of Wnt/β-catenin signaling (Waxman, 2004) that oscillates in mouse and human PSM cells (Krol et al., 2011; Matsuda et al., 2020) is also expanded and elevated in *MZpnrc2* mutants (Fig. 2M, M’ vs N, N’, RT-qPCR in O). Expression of *spry1* and *spry4*, genes encoding Fgf signaling antagonists whose orthologs oscillate in mouse and/or human PSM cells (Matsuda et al., 2020) is also expanded and elevated (Fig. 2P, P’ vs Q, Q’ and S, S’ vs T, T’, RT-qPCR in R and U). Additional genes known to oscillate in the zebrafish PSM (Krol et al., 2011) were quantified by RT-qPCR (Fig. S2). Of the 20 currently annotated genes known to oscillate in the zebrafish PSM (Krol et al., 2011), 10 (half) are overexpressed by RNA-Seq analysis in *MZpnrc2* mutants and validated by RT-qPCR (Figs 2C, S2A-F, 3C, Table S3). The oscillatory gene *caskin1* is overexpressed by RT-qPCR (Fig S2H), but unaffected in RNA-Seq analysis of *MZpnrc2* mutants (Table S3). The oscillatory gene *her2* is overexpressed in RNA-Seq analysis of *MZpnrc2* mutants, but not by RT-qPCR (Fig. S2I). An additional 5 genes are unaffected in both RNA-Seq and RT-qPCR analysis (Fig. S2I-N, Table S3), whereas among 2 genes that are under-expressed by RNA-Seq analysis, only one, *cpe*, is validated as under-expressed by RT-qPCR (Fig. S2O-P, Table S3). Overall, results show that Pnrc2 promotes decay of mRNAs that encode many developmental regulators; however, the lack of overt segmentation or additional developmental phenotypes in *MZpnrc2* mutants suggests that Pnrc2-dependent mRNAs are regulated translationally.

### Overexpressed oscillatory gene transcripts in MZpnrc2 mutants have shortened poly(A) tails

Despite overexpression of many oscillatory and segmentation-related genes, *MZpnrc2* mutants have no overt segmentation phenotype. Discordance between mRNA and protein levels might correspond with changes in the poly(A) tail status of accumulated transcripts. To examine poly(A) (pA) tail length, we performed pA tail length (PAT) assays and find that although the predominant *her1* pA tail length is ∼45 nts in both WT and *MZpnrc2* mutants, the length distribution is reduced in *MZpnrc2* mutants, which tend to have shorter tails compared to WT embryos as revealed by electrophoretic pixel intensity profiling of PAT products (Figs 3A-B, S3A, D). Similar reduced length distributions are observed for oscillatory gene transcripts *her7*, *dlc*, and *rhov* (Figs 3A-B, S3B-C, E-J). Interestingly, PAT profiling reveals a reduction in products smaller than the predominant tail length in *MZpnrc2* mutants (Fig. 3B), suggesting that accumulated transcripts in *MZpnrc2* mutants are initially deadenylated efficiently, but further deadenylation is inefficient in the absence of Pnrc2 function.

**Fig. 3.**
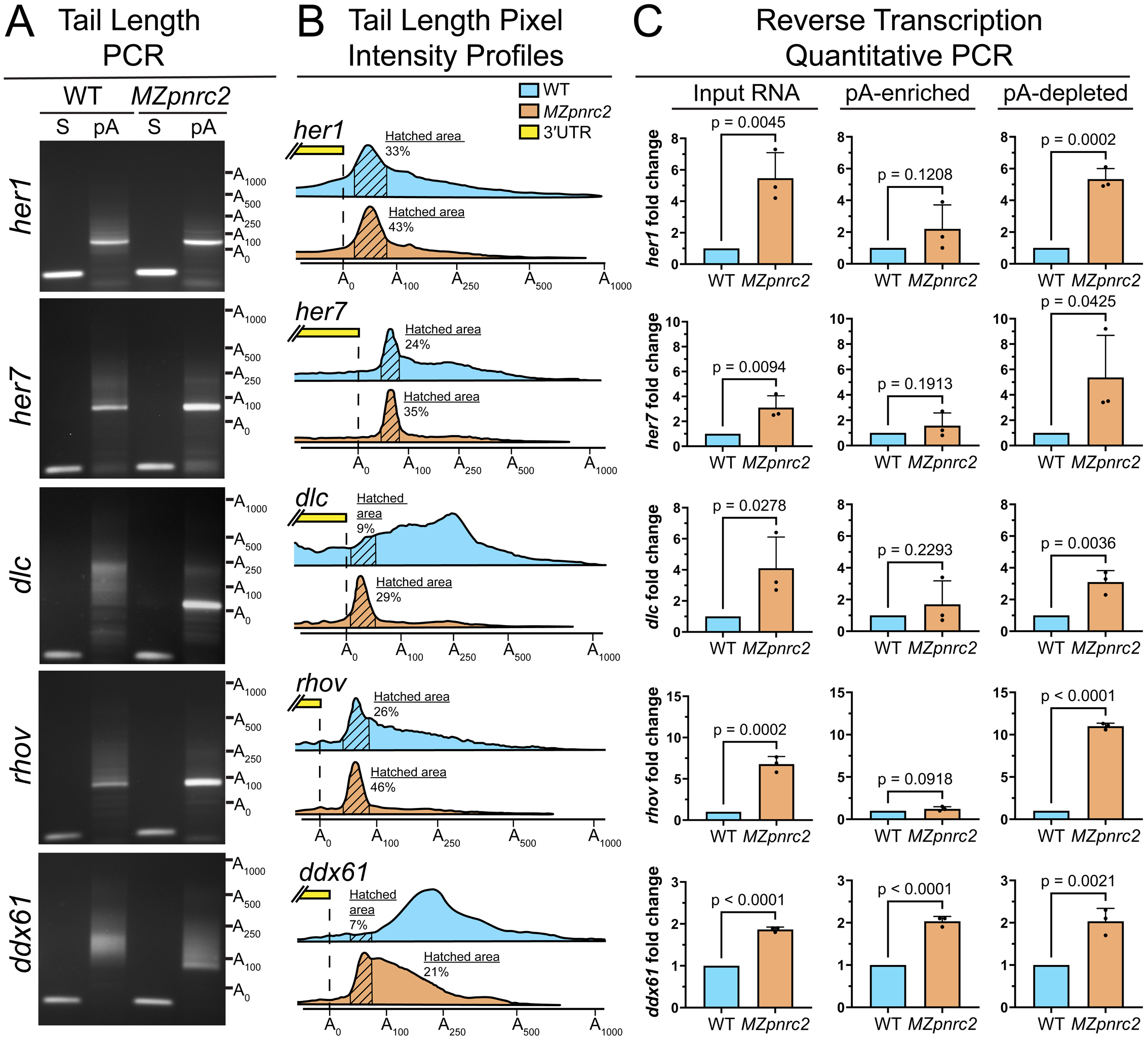
Overexpressed oscillatory gene transcripts and *ddx61* transcripts have shortened poly(A) tails in *MZpnrc2* mutants. (A) Poly(A)-tail (pA-tail) lengths of transcripts from WT and *MZpnrc2* mutants at mid-segmentation stages were measured by G/I tailing-based RT-PCR (Martin & Keller, 1998; Kusov et al., 2001), revealing a predominant pA-tail length of ∼45 nts for *her1* and ∼60 nts for *her7* and *rhov* in both WT and *MZpnrc2* mutants. A broad tail distribution pattern is observed for *dlc* in WT embryos that becomes reduced in *MZpnrc2* mutants to a predominant tail length of ∼100 nts. In contrast, *ddx61* pA-tail lengths are broadly distributed in both WT and *MZpnrc2* mutants, though the distribution is shifted to smaller tail lengths in mutants. (B) Lane profile analysis reveals that the *MZpnrc2* mutant pA-tail length distribution is reduced relative to WT for all transcripts profiled. Predominant peaks are demarcated with hatched lines (see methods), with the relative proportion of tail products in hatched regions relative to the entire plot (beginning at position A_0_) indicated. All pA-tail length results were consistent across two bio-replicates for each genotype. (C) Transcript quantification from WT and *MZpnrc2* mutant extracts at mid-segmentation stages. Total RNA (n = 50 embryos per extract per bio-replicate) was separated into pA-enriched and pA-depleted extracts by oligo-dT bead-based purification (see methods). Fold change values are presented as the mean of 3 bio- replicates with each bio-replicate value derived from the average of 2 technical replicates. Error bars show standard deviation and p-values show results from unpaired t tests. hpf = hours post fertilization; pA = poly(A) tail PCR; S = gene-specific PCR.

Because most oscillatory gene transcripts with short pA tails in *MZpnrc2* mutants would likely be transcripts that accumulate due to loss of Pnrc2 function, we quantified *her1* and additional oscillatory gene transcripts in pA-enriched and pA-depleted extracts obtained by oligo(dT) bead-based purification (see methods). The oscillatory gene transcripts *her1*, *her7*, *dlc*, and *rhov* are all significantly increased in pA-depleted extracts, but not pA-enriched extracts, derived from *MZpnrc2* mutant versus WT embryos (Fig. 3C). Transcripts with short pA tails may not be efficiently captured using oligo(dT) beads, so we quantified oscillatory gene transcripts using cDNA synthesized from total RNA extracts and primed with oligo(dT) to select for polyadenylated mRNA (rather than priming with random primers as performed for all other quantification experiments). Quantification of oligo(dT)-primed cDNA shows that oscillatory gene transcripts *her1*, *her7*, *dlc*, and *rhov* are all significantly overexpressed in *MZpnrc2* mutants (Fig. S3K-N) indicating that accumulated transcripts, although shortened, are not completely deadenylated.

### Overexpressed ddx61 transcripts in MZpnrc2 mutants are substantially polyadenylated

Accumulation of mRNA in *MZpnrc2* mutants might lead to localization to cytoplasmic granules such as processing bodies (P-bodies) that are sites of translationally repressed transcripts (Luo et al., 2018; Ostareck et al., 2014). In eukaryotic cells, P-body formation is enhanced in cells that accumulate mRNA decay intermediates (Sheth and Parker, 2003; Teixeira et al., 2005) as well as in cells treated with the translational inhibitor puromycin that results in ribosomes dissociating from mRNAs (Eulalio et al., 2007; Zheng et al., 2008).

Transcripts of several genes associated with the enriched GO term “P-body assembly” are overexpressed in *MZpnrc2* mutants (Fig. S3O-P, Table S5). These include *ddx6* and *ddx61* that are orthologous to the highly conserved *DDX6/p54* DEAD-box protein gene family that is required for P-body formation (Luo et al., 2018; Ostareck et al., 2014). Both *ddx6* and *ddx61* are expressed broadly based on single-cell RNA-Seq (Sur et al., 2023; Farrell et al., 2018). Of the two zebrafish *ddx6* ohnologs (Singh et al., 2020), *ddx61* transcript levels are 16-fold higher than *ddx6* at mid-segmentation stages (White et al., 2017), indicating that *ddx61* is the predominantly expressed ohnolog during segmentation. Upon profiling *ddx61* pA tail lengths, a broad tail length distribution is observed in both WT and *MZpnrc2* mutants, although there is a shift in distribution to shorter products in mutants (Figs 3A-B, S3E). Transcript profiling in pA-enriched and pA-depleted extracts reveals that *ddx61* is significantly increased in both fractions (Fig. 3C).

### Accumulated oscillatory gene transcripts in MZpnrc2 mutants are disengaged from translational machinery, whereas overexpressed ddx61 transcripts are translationally engaged

Because *MZpnrc2* mutants develop normally despite accumulation of numerous transcripts, we assessed engagement of *her1* and other oscillatory gene transcripts with ribosomes. We conducted polysome profiling from WT and *MZpnrc2* mutants to compare oscillatory gene transcript levels in actively translating versus non-translating fractions (see methods). The overall polysome profiles are similar in WT and *MZpnrc2* mutants (Fig. 4A), suggesting that global translation is normal in mutants; however, when comparing the relative proportion of *her1* mRNA across fractions, 63% of *her1* mRNA is found in WT polysome fractions, whereas only 27% of *her1* mRNA is found in *MZpnrc2* polysome fractions relative to total mRNA abundance across all fractions (Fig. 4B). Conversely, the proportion of *her1* mRNA in WT unbound/subunit fractions is 27%, whereas 65% of *her1* mRNA is found in *MZpnrc2* unbound/subunit fractions. The relative proportion of *her7* mRNA is similarly enriched in unbound/subunit fractions relative to polysome fractions in *MZpnrc2* mutants (Fig. 4C), as well as for *rhov* and *dlc* mRNA, although differences in mRNA proportions did not reach significance for *rhov* and *dlc* (Fig. 4D-E). Comparing oscillatory gene transcript levels between WT and *MZpnrc2* mutants reveals that *her1* transcripts in *MZpnrc2* mutants are 20-fold higher in unbound/subunit fractions, and not significantly different in monosome- and polysome-bound fractions compared to WT *her1* levels (Fig. 4G). Similar distributions were obtained for the oscillatory gene transcripts *her7*, *dlc*, and *rhov* (Fig. 4H-J). Together, these results support the hypothesis that accumulated oscillatory gene transcripts in *MZpnrc2* mutants are not efficiently translated. Because *ddx61* mRNA is increased in pA-enriched RNA extracts (Fig. 3C), we profiled *ddx61* mRNA abundance across fractions and find no differences in the proportion of *ddx61* mRNA across fractions in WT and *MZpnrc2* mutants (Fig. 4F). *ddx61* mRNA levels are 3- fold higher in *MZpnrc2* polysome fractions relative to WT polysome fractions (Fig. 4F,K), suggesting that higher engagement with ribosomes in *MZpnrc2* mutants may result in increased Ddx61 protein levels.

**Fig. 4.**
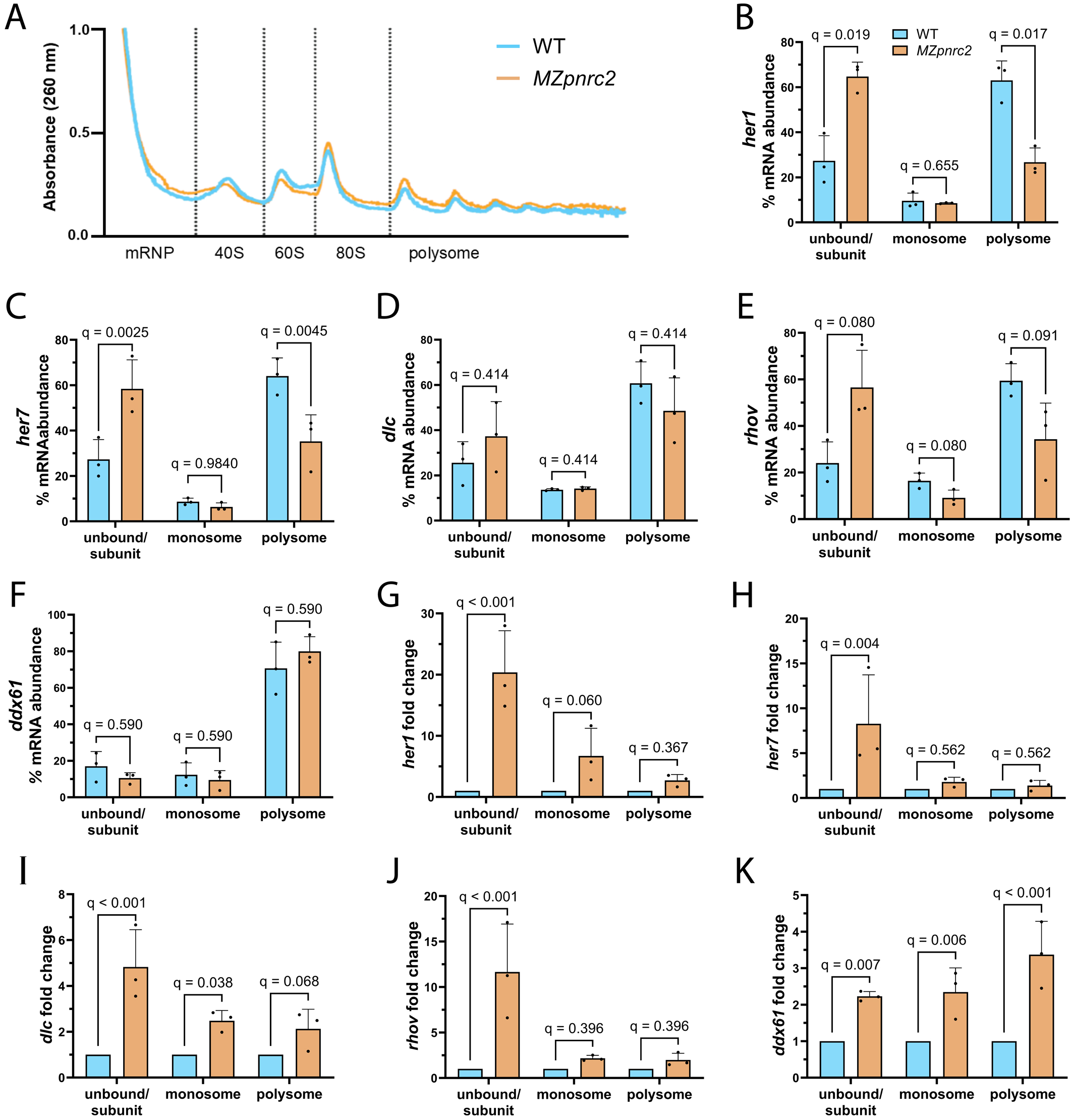
Accumulated oscillatory gene transcripts are primarily disengaged from ribosomes, whereas accumulated *ddx61* transcripts are associated with ribosomes. (A) Representative 260 nm absorbance profiles for WT and *MZpnrc2* mutant cytoplasmic extracts fractionated after cycloheximide treatment and sucrose density gradient centrifugation (n = 300 embryos per genotype at mid-segmentation stages) (see methods). Regions between dotted lines indicate fractions containing mRNAs co-sedimenting in the following groups: ribosome/subunit-unbound mRNAs (mRNP), 40S small ribosomal subunit (40S), 60S large ribosomal subunit (60S), monosome-bound mRNAs (80S), and polysome-bound mRNAs (polysome). (B-K) Transcript quantification from WT and *MZpnrc2* mutant extracts at mid-segmentation stages from unbound/subunit (combined mRNP, 40S, and 60S fractions), monosome, and polysome fractions. The proportion of transcripts in unbound/subunit, monosome, or polysome fractions relative to total transcript abundance across all fractions for WT (cyan bars) and *MZpnrc2* (orange bars) mutant samples are represented as percentages of total abundance (B-F) (see methods). Differences in transcript abundance between WT and *MZpnrc2* mutant fractions are represented as fold change (G-K) (see methods). All values are presented as the mean of 3 bio-replicates (n = 300 embryos per genotype per bio-replicate) with each bio-replicate value derived from the average of 3 technical replicates. Error bars show standard deviation and q-values show adjusted p-values from unpaired t tests adjusted with a two-stage linear step-up Benjamini, Krieger, and Yekutieli false discovery rate (FDR) procedure (Benjamini et al., 2006). nm = nanometers; mRNP = messenger ribonucleoprotein particle; S = Svedberg unit.

### Ddx6/Ddx61 protein expression and Ddx6/Ddx61-containing puncta are increased in *MZpnrc2* mutants

To corroborate *ddx61* polysome profiling results, we quantified protein levels in WT and *MZpnrc2* mutant extracts using an antibody that recognizes both Ddx6 and Ddx61. Antibodies that distinguish the two highly similar proteins are unavailable. Immunoblot results show a significant increase in Ddx6/Ddx61 protein levels in *MZpnrc2* mutants at mid-segmentation stages (Fig. 5A-B), corroborating polysome profiling (Fig 4K). To assess expression in the presomitic mesoderm (PSM), Ddx6/Ddx61 immunolabelling was performed on mid-segmentation stage embryos followed by quantitative confocal microscopy (see methods). We compared Ddx6/Ddx61 expression in maternal *pnrc2* (*Mpnrc2*) mutant embryos (that retain zygotic *pnrc2* function) to sibling *MZpnrc2* mutant embryos (that lack both zygotic and maternal *pnrc2* function) derived from the same cross to ensure similar genetic backgrounds. *Mpnrc2* mutant embryos have normal levels of oscillatory gene transcripts and develop normally (Tietz and Gallagher et al., 2020). We find that *MZpnrc2* mutants have increased Ddx6/Ddx61 expression in the PSM relative to *Mpnrc2* sibling controls (Fig 5C-E). Quantification of Ddx6/Ddx61 protein levels by measuring mean pixel intensity across the PSM reveals significantly higher pixel intensities in *MZpnrc2* mutants (Fig. 5F-H, S4A-C, G-I, M-AD).

**Fig. 5.**
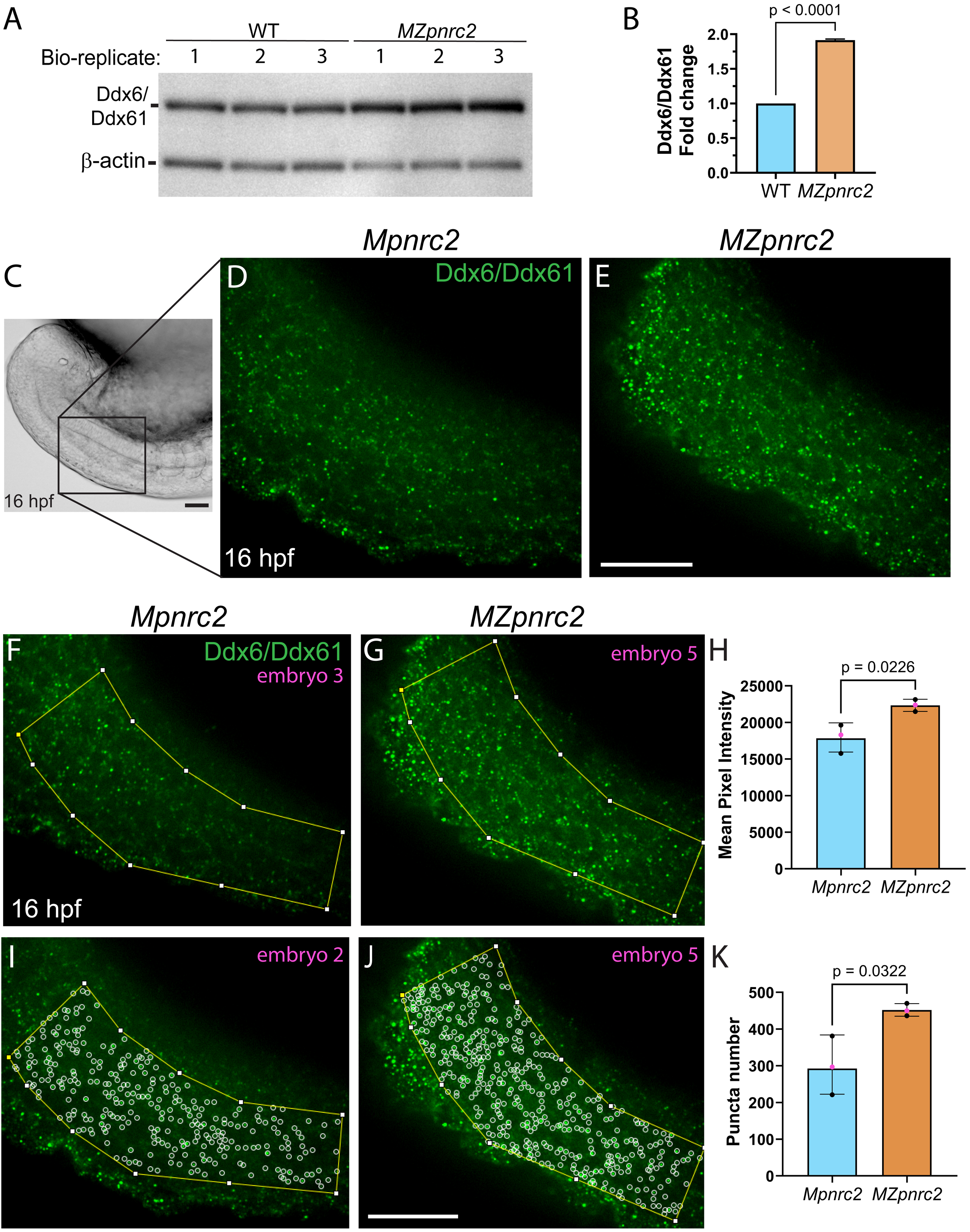
Ddx6/Ddx61 protein levels and Ddx6/Ddx61-containing puncta are increased in *MZpnrc2* mutant embryos. (A) Immunoblot showing Ddx6/Ddx61 protein levels in extracts from WT and *MZpnrc2* mutants with β-actin as a loading control (1 embryo equivalent loaded per lane at mid-segmentation stages). (B) Bar graph showing relative Ddx6/Ddx61 protein levels in WT and *MZpnrc2* mutant extracts calculated using pixel intensities from lane profile analysis of immunoblot in A. Signal intensities normalized to β-actin. Fold change value is presented as the mean of 3 bio- replicates. Error bar shows standard deviation and p-value was calculated from an unpaired t-test. (C) Brightfield image of a mid-segmentation stage embryo at 5X magnification with a box that demarcates a region containing the presomitic mesoderm (PSM) and indicates the viewable area captured by confocal microscopy at 60X magnification in Figs 5 and S4. (D-E) Ddx6/Ddx61 protein expression appears increased in representative immunolabelled *MZpnrc2* mutant embryos relative to heterozygous *Mpnrc2* sibling controls. (F-H) Quantification of Ddx6/Ddx61 protein levels by measuring mean pixel intensity in a uniformly defined region of interest (ROI) that includes the PSM (see methods). Images with mean signal intensities closest to the mean for each genotype are shown in F-G with quantification in H. (I-K) Quantification of Ddx6/Ddx61- containing puncta (see methods). Images with puncta numbers closest to the mean for each genotype are shown in I-J with quantification in K. All quantifications were performed using three embryos per genotype. Error bars show standard deviation. p-values were calculated from an unpaired t-test (H, K). Embryo numbers (magenta text in F-G & I-J) indicate embryos with quantifications closest to the mean and correspond with magenta dots indicated in bar graphs (H, K). Images for all embryos per genotype analyzed are shown in Fig. S4 S-X’. hpf = hours post-fertilization; scale bars = 50 um.

Quantification of Ddx6/Ddx61-containing puncta is also increased in *MZpnrc2* mutants (Fig. 5I- K, S4D-F, J-L, M’-AD’). Overall, our results show that *ddx61* gene products, unlike oscillatory genes, are overexpressed at both mRNA and protein levels in *MZpnrc2* mutants. Ddx6/Ddx61 expression is cytoplasmically enriched in the PSM of WT and *MZpnrc2* mutant embryos at mid-segmentation stages (Fig. 6A-C’), consistent with a role for Ddx6/Ddx61 factors in cytoplasmic post-transcriptional regulation.

**Fig. 6.**
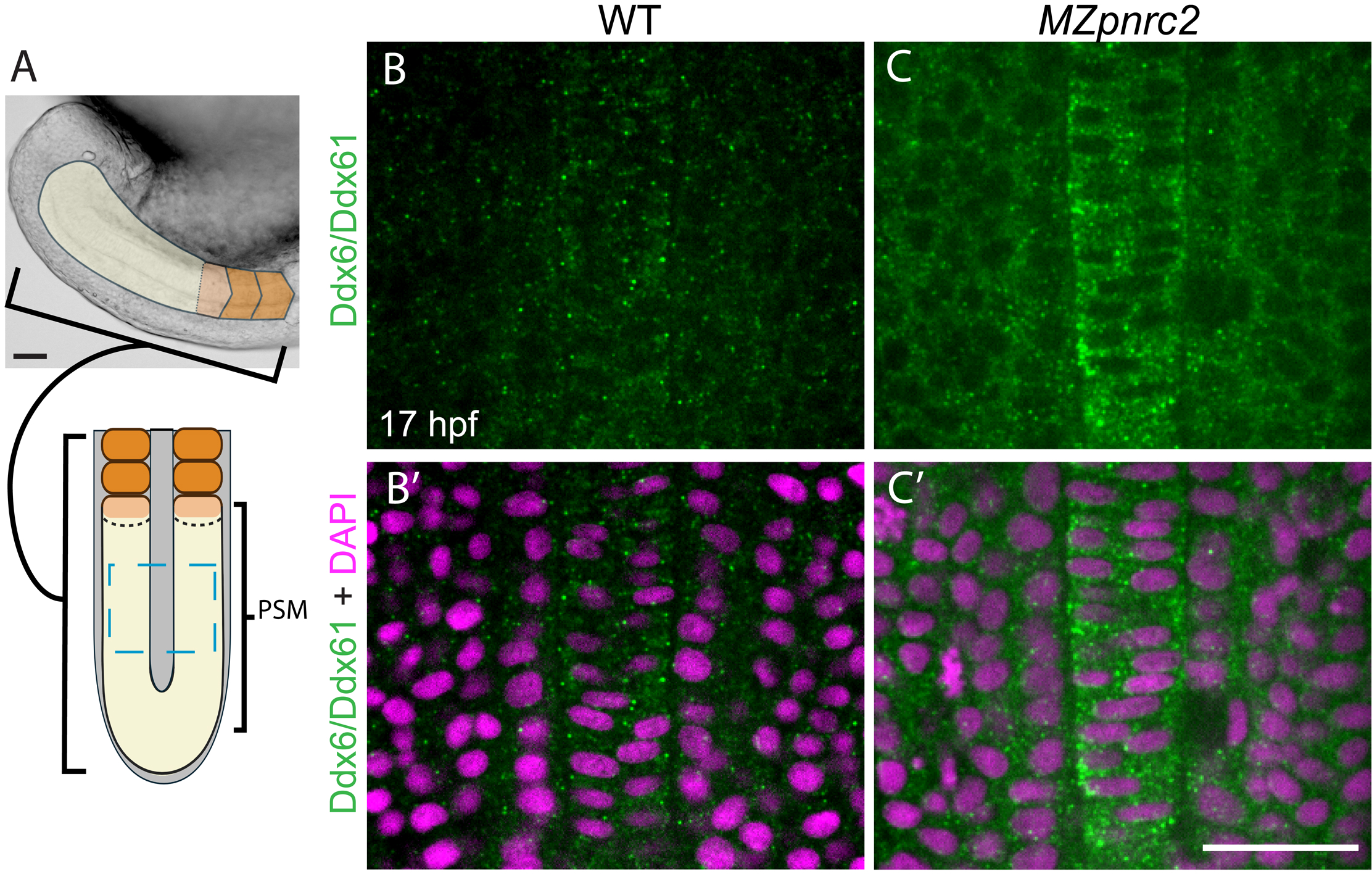
Ddx6/Ddx61 expression is cytoplasmically-enriched in the presomitic mesoderm. (A) Brightfield image of a mid-segmentation stage embryo (from Fig. 5C) with overlayed illustration to highlight the presomitic mesoderm (PSM) in yellow and formed somites in orange. A newly forming somite is shown in light orange with a hatched posterior boundary. Rotated schematic illustrates a dorso-posterior view (anterior to the top) with a hatched blue box showing the viewable area captured by confocal microscopy in B-C. (B-C’) Ddx6/Ddx61 protein expression is cytoplasmically enriched in WT and *MZpnrc2* mutant embryos. Representative images are shown with DAPI counterstaining in B’-C’ (n = 5 embryos per genotype analyzed). hpf = hours post-fertilization; scale bars = 50 um.

### MZpnrc2 mutants are sensitized to deadenylation inhibition

Broad ectopic expression of a CCR4-NOT dominant negative deadenylase via mRNA injection into zebrafish embryos has previously been shown to disrupt segmentation and prevent pA tail shortening of oscillatory gene transcripts *her1* and *her7* (Fujino et al., 2018). Consistent with published work, we show that *Myc-Cnot7DN* mRNA injection causes segmentation defects in WT embryos, with over 80% of injectants exhibiting completely malformed segments (Fig. S5A-D). Because segmentation clock gene transcripts accumulate in *MZpnrc2* mutants, we hypothesized that *MZpnrc2* mutants would be sensitized to deadenylation inhibition. To test this idea, we pursued an inducible deadenylase inhibitor expression strategy by injecting a plasmid containing a heat-shock inducible version of the deadenylase inhibitor.

After inducing expression of Myc-tagged Cnot7DN for 30 minutes at mid-segmentation stages, WT and *MZpnrc2* mutants were scored for segmentation defects 6 hours post-heat shock (see methods). Mosaic expression of Myc-Cnot7DN leads to abnormal segment formation in *MZpnrc2* mutants, but not in WT embryos (Fig. 7A-E), consistent with the hypothesis that *MZpnrc2* mutants are sensitized to deadenylase inhibition. Using anti-Myc immunohistochemistry on plasmid-injected embryos, we confirmed that Myc-Cnot7DN expression was induced post-heat shock and was absent in no-heat shock controls (Fig. 7F-G). No significant Myc-Cnot7DN expression pattern differences were observed between genotypes (Fig. S5I-M), showing that *MZpnrc2* mutant sensitivity is not due to differences in transgene induction. When comparing transgene expression patterns between *MZpnrc2* mutants with and without somite defects, all *MZpnrc2* mutants with very high Myc-Cnot7DN expression (i.e. embryos showing broad expression with little to no mosaicism) displayed somite defects, though overall, expression pattern differences between *MZpnrc2* mutants with and without somite defects did not reach significance (p = 0.056) (Fig. S5N).

**Fig. 7.**
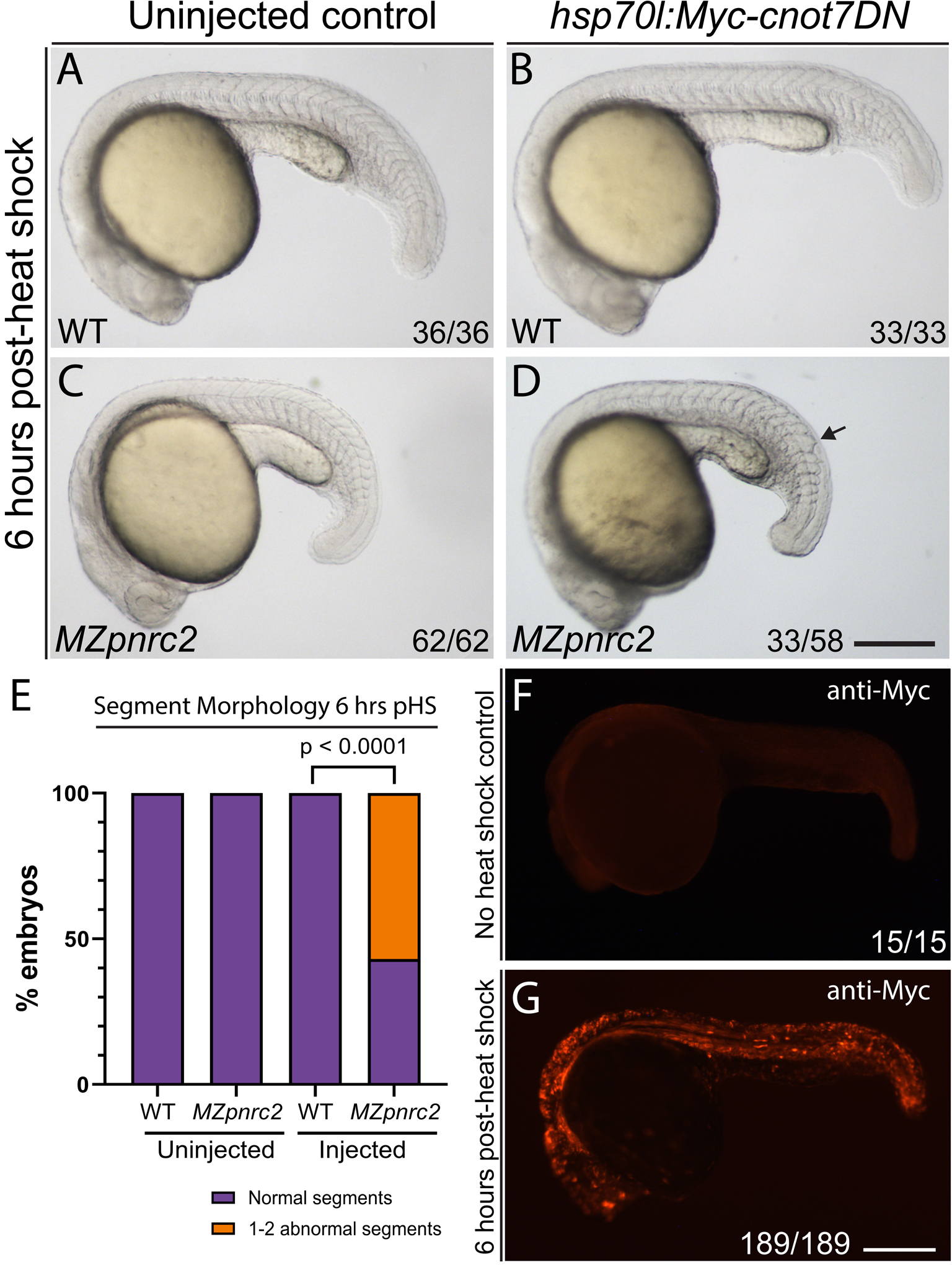
Induced expression of a dominant-negative deadenylase has no effect in WT embryos but induces segmentation defects in *MZpnrc2* mutants. WT and *MZpnrc2* mutants were injected with 20 pgs *hsp70l:Myc-cnot7DN* plasmid at the 1-cell stage and raised alongside uninjected clutch-mate controls to ∼14 hpf. Injected and uninjected controls were then heat shocked at 37 C for 30 minutes and raised at standard temperature (28.5 C) for 6 hours post-heat shock (pHS) with morphological assessment at hourly intervals (see methods). (A-D) Representative embryos are shown for (A) uninjected WT embryos (n = 36), (B) *hsp70l:Myc- cnot7DN*-injected WT embryos (n = 33), (C) uninjected *MZpnrc2* mutants (n = 62), and (D) *hsp70:Myc-cnot7DN*-injected *MZpnrc2* mutants (n = 58). Arrow in D indicates an abnormal segment. (E) Bar graph shows the proportion of embryos with normal and/or abnormal segments, indicating that injected *MZpnrc2* mutants are the only group shown in A-D with somite defects. (F-G) A subset of injected embryos was reserved as no heat-shock controls and were negative for Myc-Cnot7DN expression by anti-Myc immunohistochemistry (n = 15) (F), whereas mosaic expression was observed in injected embryos 6 hours pHS (n = 189) (G) (representative images shown in F-G). P-value was calculated using a Fisher’s Exact test comparing the number of embryos with or without segment defects. Plasmid injection followed by heat-shock induction was performed twice independently with consistent results. hpf = hours post fertilization; pHS = post-heat shock; DN = dominant negative; scale bar = 300 um.

### MZpnrc2 mutants are sensitized to depletion of Ddx61 and Ddx6 function

Because *ddx61* is overexpressed at both mRNA and protein levels, we hypothesized that depleting Ddx61 function in *MZpnrc2* mutants might lead to segmentation and/or additional phenotypes. Using a highly efficient F0 CRISPR knockdown strategy (Hoshijima et al., 2019; Klatt Shaw and Mokalled, 2021), we find that single *ddx61* crispants and single *ddx6* crispants develop normally up to 5 days post-fertilization (dpf) and can be raised without significant losses up to adulthood (data not shown), consistent with *ddx6* and *ddx61* genetic mutant analyses (Pronobis et al., 2025). To estimate F0 CRISPR efficiency (Fig. S6A), we employed high resolution melt analysis (HRMA) to detect CRISPR-induced mutations and confirmed that all CRISPR-injected embryos were mutagenized (Fig. S6B-C). Because *ddx6* might compensate for loss of *ddx61* function, we also targeted *ddx6*. In *ddx6*;*ddx61* crispants, *ddx6* and *ddx61* mRNA levels are more than 50% reduced at 18 hpf compared to uninjected sibling controls (Fig. S6D-E), likely because the premature termination codon-containing transcripts created by our targeting strategy are degraded by NMD. Ddx6/Ddx61 protein levels are also reduced in *ddx6*;*ddx61* crispants at 18 hpf and are nearly absent by 4 dpf (Fig. S6F). To assess whether *MZpnrc2* mutants are sensitized to Ddx6/Ddx61 depletion, we compared *ddx6*;*ddx61* crispant phenotypes in WT and *MZpnrc2* mutant backgrounds over time. At mid-segmentation stages, most *ddx6;ddx61* crispants are morphologically normal in WT and *MZpnrc2* mutant backgrounds (Fig. 8A-B, E-F, I). By 19 hpf, ∼30% of *MZpnrc2 ddx6*;*ddx61* crispants exhibit morphological abnormalities, whereas *ddx6*;*ddx61* crispants in the WT background (hereafter referred to as *ddx6*;*ddx61* crispants) as well as uninjected controls for both genotypes are predominantly normal (Fig. 8C-D, G-H, J). Defects worsen with time in *MZpnrc2 ddx6*;*ddx61* crispants, with nearly all crispants showing abnormalities at 60 hpf when *ddx6*;*ddx61* crispants are mostly normal (Fig. S7A-J). By 84 hpf, nearly half of *ddx6*;*ddx61* crispants show abnormal phenotypes similar to those observed earlier in *MZpnrc2 ddx6;ddx61* crispants (Figs 9A-D, S7K). Similar proportions of abnormal phenotypes are observed at 112 hpf and 120 hpf (Figs 9E-F, S7L) and correspond with strongly reduced Ddx61/Ddx6 protein levels in *ddx6*;*ddx61* crispants (Figs 9G, S6F-G). Overall, these results support that *MZpnrc2* mutants are sensitized to reduced Ddx6/Ddx61 function.

**Fig. 8.**
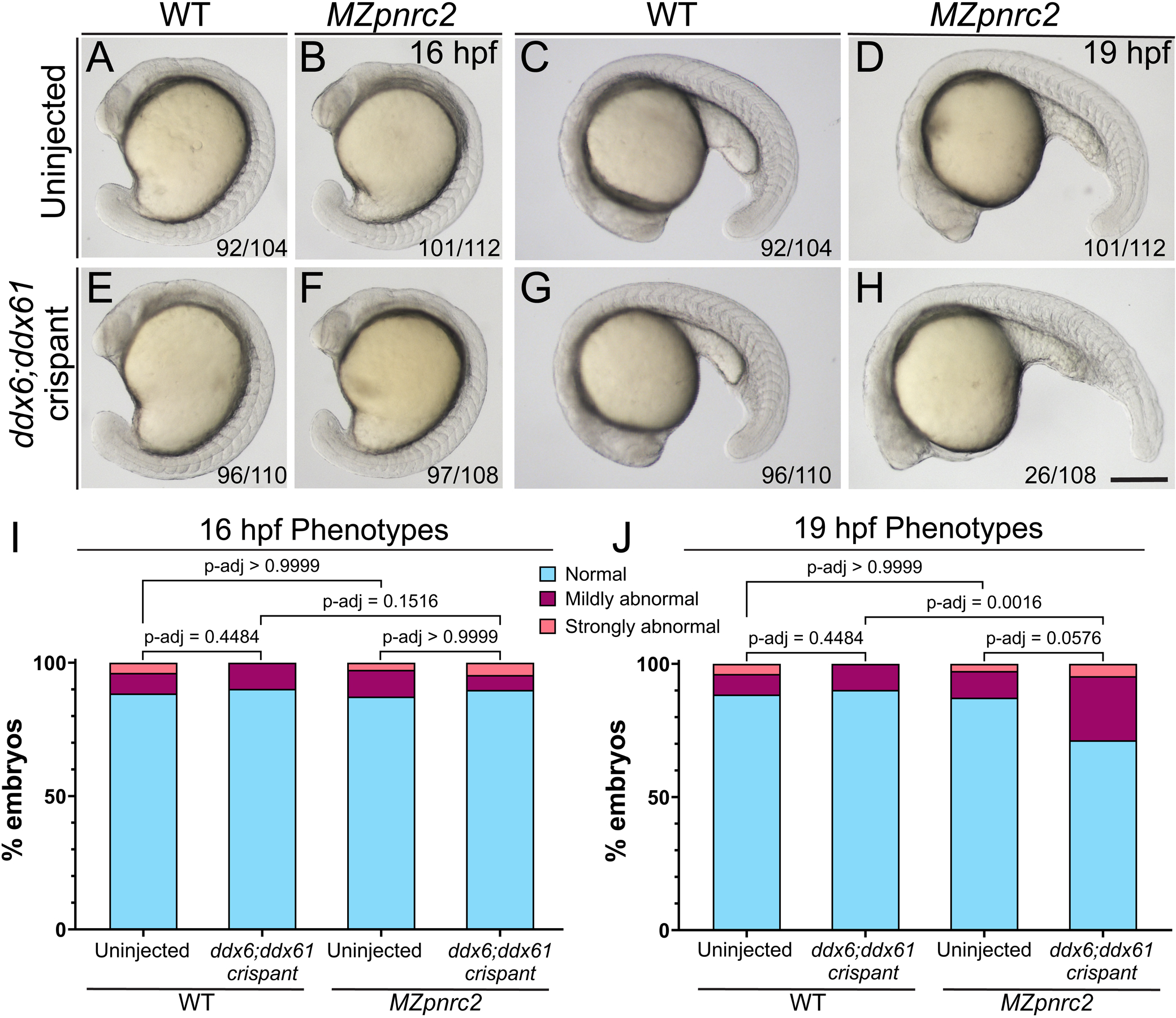
*MZpnrc2* mutant embryos with depleted Ddx6 and Ddx61 function first show abnormal phenotypes at late segmentation stages. (A-H) Representative embryos are shown at 16 hpf and 19 hpf, respectively, for (A, C) uninjected WT and (B, D) uninjected *MZpnrc2* mutants, (E, G) *ddx6*;*ddx61* crispants (on a WT background), (F, H) *MZpnrc2 ddx6*:*ddx61* crispants. (I-J) Bar graphs show the proportion of embryos with normal and abnormal phenotypes at 16 hpf (I) and 19 hpf (J). Adjusted p-values were calculated using a Fisher’s Exact test comparing the number of embryos within each phenotype class (normal., mildly abnormal., strongly abnormal) and corrected for multiple pairwise comparisons using a Bonferroni adjustment (Bonferroni, 1935; Bonferroni, 1936; Dunn, 1961). Four knockdown experiments were independently performed and produced consistent results. S = somite stage; scale bar = 250 um.

**Fig. 9.**
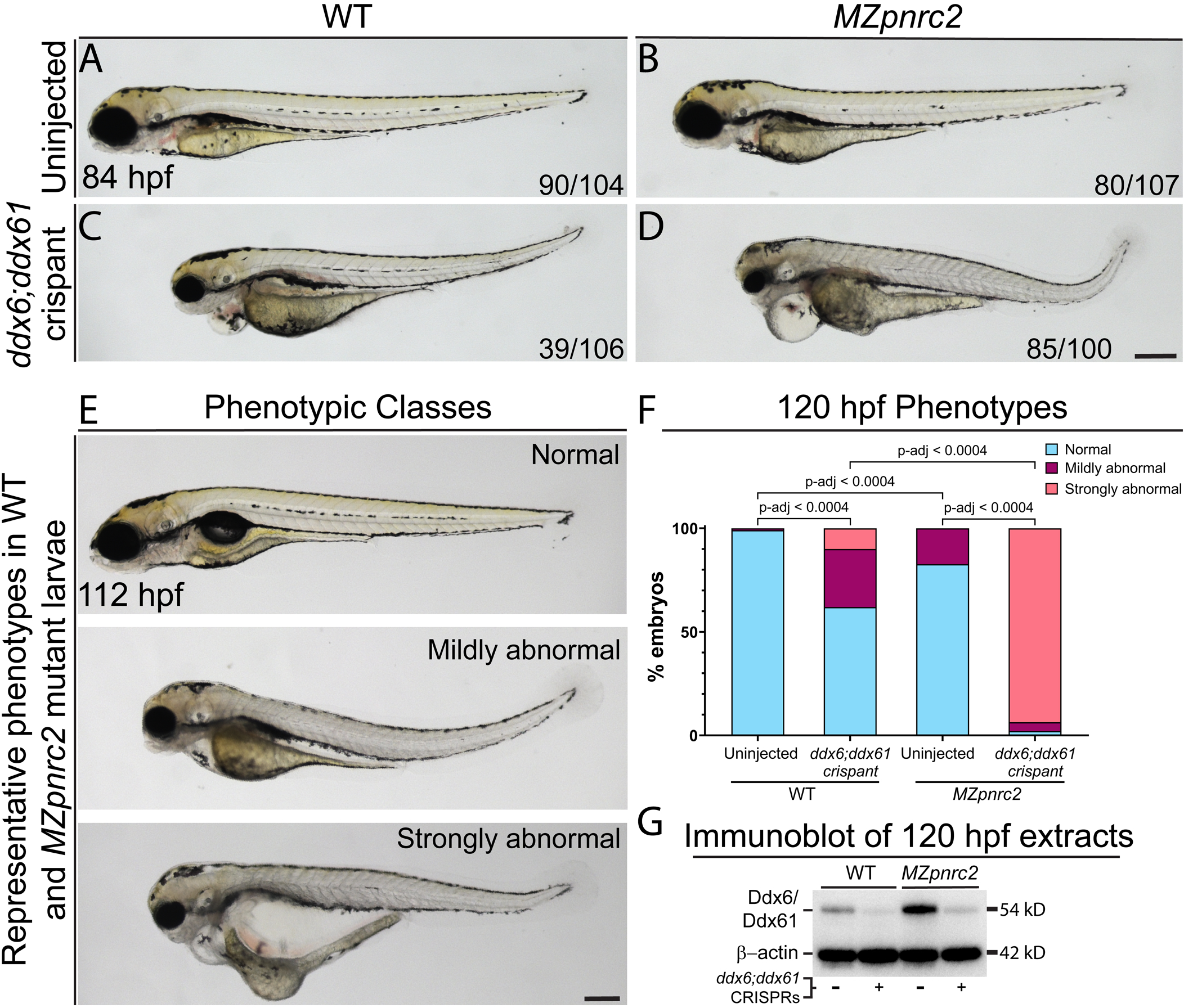
*MZpnrc2* mutant larvae are sensitized to depletion of Ddx6 and Ddx61. (A-D) Representative images are shown at 84 hpf for (A) uninjected WT, (B) uninjected *MZpnrc2*, (C) *ddx6*;*ddx61* crispants (on a WT background), and (D) *MZpnrc2 ddx6*:*ddx61* crispants. (E) By 112 hpf, *ddx6;ddx61* crispants exhibit mild to severe abnormalities that occur at higher frequency in *MZpnrc2* mutants (representative phenotypic classes shown in E with phenotypic proportions in Fig. S6L). (F) Bar graph showing proportion of larvae at 120 hpf with normal and abnormal phenotypes, indicating a synergistic effect from loss of *pnrc2*, *ddx6*, and *ddx61*. Proportions were calculated from the following groups: uninjected WT, n = 112; *ddx6;ddx61* crispant, n = 100; uninjected *MZpnrc2*, n = 81; *MZpnrc2 ddx6;ddx61* crispant, n = 47. Adjusted p-values were calculated using a Fisher’s Exact test comparing the number of embryos within each phenotype class (normal., mildly abnormal., strongly abnormal) and corrected for multiple pairwise comparisons using a Bonferroni adjustment (Bonferroni, 1935; Bonferroni, 1936; Dunn, 1961). Four independent knockdown experiments were performed and produced consistent results. (G) Immunoblot showing increased Ddx6/Ddx61 protein levels in *MZpnrc2* mutants and efficient depletion of Ddx61/Ddx6 protein in *ddx6;ddx61* crispants at 120 hpf. One-half larval equivalents were loaded in each lane with β-actin shown as a loading control. Immunoblot results were consistent across two independent experiments. hpf = hours post-fertilization; dpf = days post-fertilization; scale bars = 250 um.

To determine whether *her1* expression is affected in *ddx6;ddx61* crispants, we performed *her1* in situ hybridization at mid-segmentation stages. Although the *her1* expression pattern in *ddx6*;*ddx61* crispants was indistinguishable from uninjected WT controls (Fig. 10A,A’ vs B,B’), *MZpnrc2 ddx6*;*ddx61* crispants showed an expanded misexpression phenotype compared to control *MZpnrc2* mutants (Fig. 10C,C’ vs D, D’). We quantified *her1* levels by RT-qPCR and find that *her1* is more than 1.5-fold increased in *MZpnrc2 ddx6*;*ddx61* crispants relative to control *MZpnrc2* mutants (Fig. 10E), but is unaffected in *ddx6*;*ddx61* crispants. In contrast, *her7* expression is reduced, whereas *dlc* and *rhov* expression is unaffected in *MZpnrc2 ddx6*;*ddx61* crispants (Fig. 10F-H). Overall, normal levels of oscillatory gene transcripts in *ddx6*;*ddx61* crispants suggest that Ddx61 and Ddx6 are not required for mRNA decay when Pnrc2 is functional. The synergistic effect of loss of *pnrc2*, *ddx6*, and *ddx61* shows that Ddx6 and Ddx61 partially compensate for loss of Pnrc2 function.

**Fig. 10.**
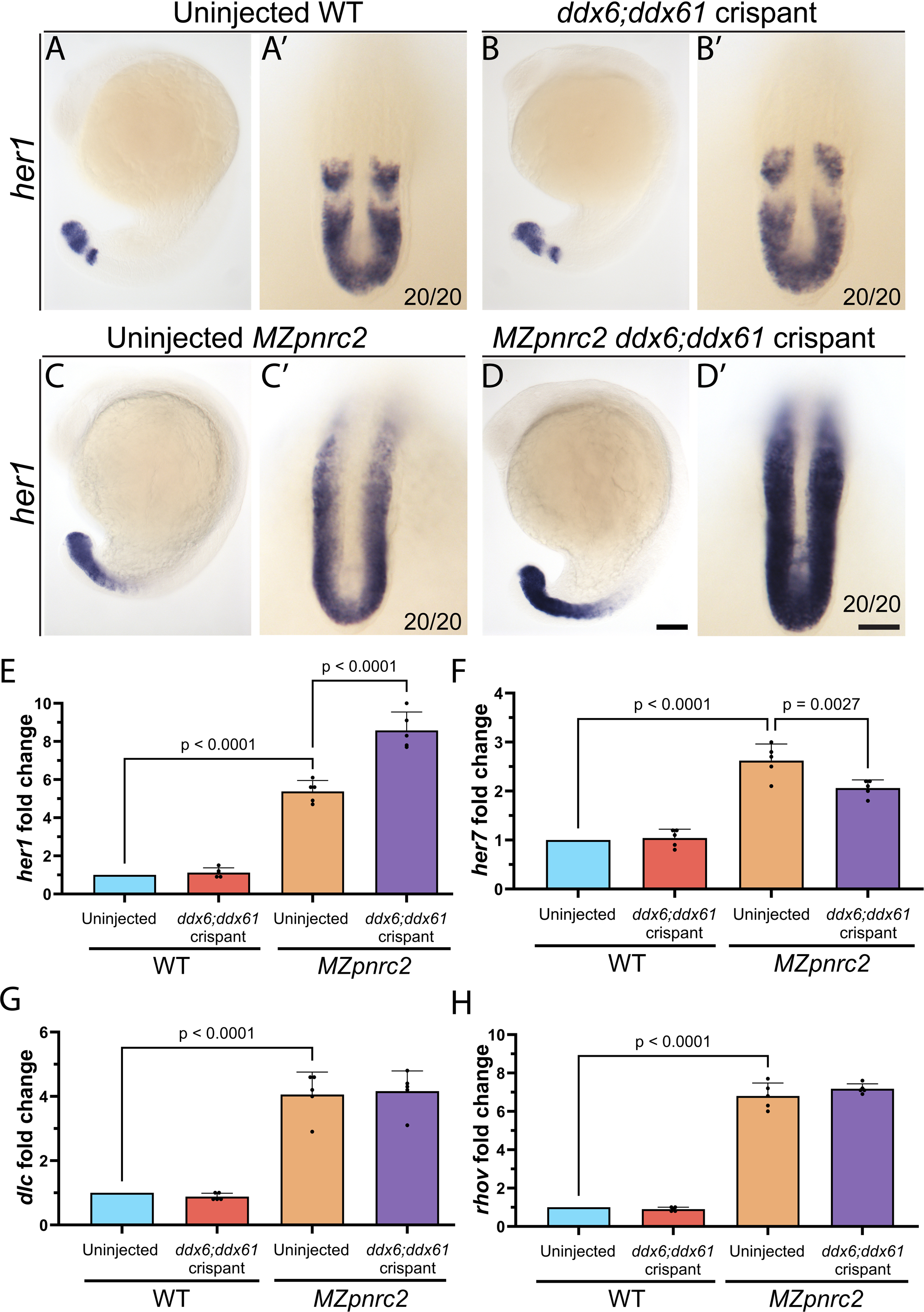
Injection of *ddx6;ddx61* CRISPRs does not affect *her1* mRNA levels in WT embryos, but increases *her1* accumulation in *MZpnrc2* mutants. WT and *MZpnrc2* mutants were injected with *ddx6;ddx61* CRISPRs and raised to 18 hpf alongside uninjected clutch-mates as controls. Uninjected and injected embryos were probed for *her1* expression by in situ hybridization (n = 20 embryos per genotype per condition) and oscillatory gene transcript levels were quantified by RT-qPCR (n = 10 embryos per genotype per condition; 5 bio-replicates per condition with bio-replicate values derived from the average of 2 technical replicates each). (A-D’) Although *her1* expression in *ddx6;ddx61* crispants is indistinguishable from uninjected clutch-mates (A, A’ vs B, B’), *her1* expression appears expanded in *MZpnrc2 ddx6;ddx61* crispants relative to uninjected *MZpnrc2* mutant clutch-mates (C, C’ vs D, D’). Lateral views are shown in A-D, magnified dorsal views are shown in A’-D’. (E-H) Bar graphs showing RT-qPCR results with fold change values derived from the mean of 5 bio-replicates. Out of four quantified oscillatory gene transcripts, only *her1* expression is significantly increased in *MZpnrc2 ddx6;ddx61* crispants (E vs F-H). Oscillatory gene transcript expression is unaffected in *ddx6;ddx61* crispants on the WT background (E-H). Adjusted p-values calculated using a one-way ANOVA with a Dunn- Sidák correction for multiple comparisons (Sidák,1967; Abdi, 2007). Scale bars = 100 um (A-D); dorsal view = 50 um (A’-D’).

## Discussion

In this work, we show that overexpressed transcripts in *MZpnrc2* mutants are primarily protein-coding transcripts, whose sequences are enriched for features known to promote transcript decay, including uORFs, long 3′UTRs, and mRNA destabilization sequences. Human PNRC2 bridges NMD machinery by directly binding NMD factors Upf1 and Dcp1 (Cho et al., 2009; Lai et al., 2012). Although depleting *upf1* via morpholino (MO) injection in zebrafish embryos has no effect on *her1* expression, co-depletion of *upf1* and *pnrc2* via MO injection enhances *her1* misexpression compared to depletion of *pnrc2* alone (Gallagher et al., 2017), suggesting that Upf1 is important for the decay of *her1* and additional overexpressed transcripts in *MZpnrc2* mutants.

Additional features of overexpressed transcripts in *MZpnrc2* mutants include enrichment of mRNA destabilizing sequences, including PREs and AREs that are present in the oscillatory gene transcripts *her1*, *her7*, and *dlc*. Among overexpressed transcripts, many correspond to genes that are dynamically regulated, including oscillatory and/or developmental genes that function in developmental signaling pathways. Remarkably, despite overexpression of >1,700 transcripts, segmentation and other developmental processes appear normal in *MZpnrc2* mutants, although many fail to thrive as juveniles and adults. We previously suggested that absence of segmentation defects is likely because accumulated mRNAs in *MZpnrc2* mutants are not efficiently translated (Gallagher et al., 2017), though the mechanism was unknown. In the current study, we explored mechanisms driving discordant expression of mRNA and protein in *MZpnrc2* mutants by assessing the polyadenylation status of *her1* and other transcripts, their level of engagement with translational machinery, and the impact of P-body factors Ddx6 and Ddx61 on the *MZpnrc2* overexpression phenotype.

### MZpnrc2 mutants have shortened poly(A) tails and are sensitized to deadenylation inhibition

Poly(A)-tail length analysis suggests that many accumulated oscillatory gene transcripts have reduced tail lengths in *MZpnrc2* mutants, but the effect this would have on translation of accumulated oscillatory gene transcripts in *MZpnrc2* mutants remains unknown. In early zebrafish embryos, tail lengths are strongly coupled to translation efficiency (TE), but this coupling is reduced by gastrulation whereby poly(A) tail shortening primarily leads to mRNA destabilization with little effect on TE (Subtelney et al., 2014). Furthermore, in many eukaryotic model species analyzed, highly-expressed transcripts have short tails (Lima et al., 2018), indicating that short tails alone are not sufficient to repress translation. Therefore, the observations that accumulated oscillatory gene transcripts in *MZpnrc2* mutants have shortened tails and that most accumulated transcripts are not associated with ribosomes are potentially not mechanistically connected. Although the relationship between poly(A) tail length and TE are not directly explored in the current study, the finding that *MZpnrc2* mutants are sensitized to deadenylation inhibition during segmentation demonstrates the importance of deadenylation function when mRNA decay is dysfunctional.

### Depletion of RNA binding proteins Ddx6 and Ddx61 leads to developmental defects in MZpnrc2 mutants

We show here that Ddx6/Ddx61 RNA binding proteins are overexpressed at both mRNA and protein levels in *MZpnrc2* mutants, and that Ddx6/Ddx61-containing puncta, which likely represent P-bodies, are increased in the presomitic mesoderm. Depletion of *ddx6* and *ddx61* function in *MZpnrc2* mutants increases *her1* accumulation and leads to strong morphological phenotypes that are observed at much lower frequency in *ddx6;ddx61* crispant controls, showing that *MZpnrc2* mutants are sensitized to loss of *ddx6* and *ddx61* function. Future work will assess whether Pnrc2-regulated transcripts are over-translated upon depletion of Ddx6 and Ddx61 function in *MZpnrc2* crispants. It is possible that Ddx6/ Ddx61 promotes tail-shortening of accumulated mRNAs in *MZpnrc2* mutants, however a connection between deadenylation inhibition and Ddx6/Ddx61 depletion experiments has not been established.

Morphological defects are not observed in *ddx6*;*ddx61* crispants at 1 dpf despite substantial Ddx6/Ddx61 protein depletion, potentially because maternally provided *ddx6/ddx61* function is sufficient for normal early development. The absence of early phenotypes differs from previous work that showed that *ddx6* and *ddx61* splice-blocking morpholino (MO) injection leads to a curved tail phenotype in 24 hpf embryos (Zampedri et al., 2016), which could be due to differences in knockdown approach and/or efficiency. Despite early differences, embryos injected with *ddx6* and *ddx61* MOs survive to at least 5 dpf (Zampedri et al., 2016), similar to what we observed for *ddx6*;*ddx61* crispants. More recently, genetic analysis of *ddx6* and *ddx61* mutants showed that doubly homozygous *ddx6;ddx61* mutants are inviable by 1 dpf (Pronobis et al., 2025). Although doubly homozygous *ddx6*;*ddx61* genetic mutants also have maternally provided *ddx6/ddx61* function, we suspect that the differences between doubly homozygous *ddx6*;*ddx61* genetic mutants versus *ddx6*;*ddx61* crispants and morphants is likely because genetic mutants completely lack zygotic Ddx6/Ddx61 protein whereas crispants and morphants likely have low levels of zygotic Ddx6/Ddx61 protein. Further analyses of genetic mutants will clarify the role of Ddx6/Ddx61 function during early development.

In contrast to *ddx6*;*ddx61* crispants that appear largely normal up to 60 hpf, *MZpnrc2 ddx6;ddx61* crispants show abnormalities as early as 19 hpf. We hypothesize that *MZpnrc2* mutants require more Ddx6/Ddx61 function than WT embryos because of differences in transcript accumulation. In fact, Ddx6/Ddx61 protein levels may be increased in *MZpnrc2* mutants as a compensatory response that buffers against the negative effects of transcript accumulation. The finding that *her1* mRNA levels in *MZpnrc2* mutants are further increased in *ddx6*;*ddx61* crispants suggests that P-body factors promote *her1* mRNA decay in the absence of Pnrc2. P-bodies are thought to primarily function as sites of mRNA storage rather than decay, supported by observations that mRNA decay continues in the absence of P-bodies (Decker et al., 2007; Eulalio et al., 2007; Stoecklin et al., 2006). Early reports of mRNA decay in P-bodies are attributable to artifacts of the MS2 tethering system (Garcia & Parker, 2015; Heinrich et al., 2017; Tutucci et al., 2018). It is also possible that Ddx6/Ddx61 promotes *her1* mRNA decay in a P-body-independent manner, as both proteins show diffuse cytoplasmic expression in addition to enrichment in cytoplasmic puncta in zebrafish embryos (Figs. 5-6, S4) and Zampedri et al., 2016). Future work will clarify the extent to which *her1* mRNA is degraded in the cytoplasm versus in P-bodies and the dependence of these activities on *pnrc2* function.

It is intriguing that among the 4 oscillatory gene transcripts we quantified, only *her1* levels are increased in *MZpnrc2 ddx6*;*ddx61* crispants. It is possible that the *her1* transcript contains features that promote Ddx6/Ddx61-dependent mRNA decay that are absent in *her7*, *dlc*, and *rhov* transcripts. Alternatively, the level of Ddx6/Ddx61 depletion in *MZpnrc2* mutants may not have been sufficient to detect accumulation of *her7*, *dlc*, and *rhov* oscillatory gene transcripts. Future analysis using genetic mutants will help to distinguish these possibilities.

## Conclusions

*Pnrc2* promotes rapid mRNA decay during vertebrate segmentation, including the decay of many dynamically expressed oscillatory gene transcripts. Overexpressed transcripts in *MZpnrc2* mutants are enriched for mRNA-destabilizing motifs that are associated with translational repression and mRNA decay. *MZpnrc2* mutants develop normally despite dysfunctional mRNA decay, likely because transcripts that depend on Pnrc2 for rapid decay are inefficiently translated. *MZpnrc2* mutants have shortened poly(A) tails and inhibiting deadenylation leads to somite defects in sensitized *MZpnrc2* mutants, potentially due to over-translation of Pnrc2-regulated transcripts. Upregulation of *ddx61* at both mRNA and protein levels suggests that compensatory mechanisms are employed in decay-deficient *pnrc2* mutants. Loss of *pnrc2*, *ddx6*, and *ddx61* leads to synergistic and strong developmental phenotypes, which may be due to over-translation of Pnrc2-regulated transcripts. Overall, our results show that Ddx6 and Ddx61 are required for normal embryonic development in mRNA-decay deficient *MZprnc2* mutants.

## Materials and methods

### Animal stocks and husbandry

All protocols using zebrafish were approved by the Ohio State University Institutional Animal Care and Use Committee (protocol 2012A00000113). Adult zebrafish strains (*Danio rerio*) were kept at 28.5°C on a 14 hour (h) light/10 h dark cycle and obtained by natural spawning or *in vitro* fertilization (Westerfield, 2007). Embryos and larvae were staged according to Kimmel et al.,1995. The *pnrc2^oz22^*line has been described previously (Gallagher et al., 2017).

### DNA extraction and genotyping

Individual embryos and adult fin tissue were lysed in 50 ul 1X ThermoPol Buffer (NEB) at 95°C for 10 minutes, digested at 55°C for 1 hour in 1 mg/ml Proteinase K (BP1700, ThermoFisher), followed by Proteinase K inactivation at 95°C for 10 minutes. Molecular identification of *pnrc2^oz22^*carriers by PCR amplification followed by NsiI-HF digestion to distinguish cleavable WT from uncleavable mutant amplicons was performed as previously described (Tietz et al., 2020).

CRISPR mutagenesis efficiency was analyzed by high resolution melt analysis (HRMA) on a CFX Duet instrument (Bio-Rad) using 1 ul of DNA extract as template in a 20 ul reaction with Precision Melt Mix following standard manufacturer procedures (Bio-Rad). gRNA and primer sequences are listed in Table S6. Melt analysis was performed using Precision Melt Analysis Software v1.2 (Bio-Rad).

### RNA-Seq sample collection and sequencing

Whole embryos at mid-segmentation stages (n = 50 embryos per biological replicate for each genotype) were solubilized in Trizol for RNA extraction and purified following standard procedures (Thermo Fisher). All genotypes were prepared in biological triplicate. 2 ug of total RNA from each sample was processed at Novogene using a whole transcriptome sequencing workflow with ribosomal RNA (rRNA) depletion that enables sequencing of RNA irrespective of polyadenylation status. Each sample was depleted of rRNA using a Ribo-Zero Kit (Illumina), followed by chemical fragmentation and 150 bp paired-end library synthesis using a NEBNext Ultra II Directional RNA kit (NEB). Libraries were sequenced on the Illumina NovaSeq 6000 platform.

### RNA-Seq data analysis

High-throughput sequencing reads were trimmed to remove adapter and primer sequences using Cutadapt version 3.4 (Martin, 2011) and pseudoaligned to the zebrafish GRCz11/danRer11 transcriptome reference using Kallisto (version 0.46.2), an RNA-Seq tool that rapidly quantifies transcript abundance without the need for individual base alignment (Bray et al., 2016). Cutadapt and Kallisto were installed and run in Python (version 2.7.5) using standard procedures described in online guides available at https://cutadapt.readthedocs.io/en/stable/guide.html and https://pachterlab.github.io/kallisto/manual. Prior to running Kallisto, all sequence data files were assessed using the quality control tool FastQC version 0.12.1 (Andrews, 2010) and no quality issues were identified in relevant modules for RNA-Seq analysis (data not shown). Transcripts with <5 estimated read counts in at least one sample were filtered out to reduce noise.

Differential gene expression analysis was performed using Sleuth version 0.30.1, a program that uses variance decomposition to identify biological differences between RNA-Seq samples (Pimentel et al., 2017). Sleuth was run in RStudio version 2023.06.2 Build 561 (Posit Software) using standard procedures described in an online guide at https://pachterlab.github.io/sleuth_walkthroughs/trapnell/analysis.html. Batch effects including effects due to sequencing lane or biological replicate differences were tested by conducting differential gene expression analysis using Sleuth version 0.30.1 to compare samples run on different lanes (i.e., lane 1 vs lane 2) (Table S1) or samples obtained from different biological replicates independent of genotype, i.e., biological replicates 1 (WT & *MZpnrc2*) vs biological replicates 2 (WT & *MZpnrc2*) vs biological replicates 3 (WT and *MZpnrc2*) (Table S4). There were no differentially expressed transcripts detected when comparing expression from samples run on different sequencing lanes nor were there differences in expression detected when comparing samples from different biological replicates that were independent of genotype (data not shown). A summary of the RNA-Seq processed data, Kallisto normalized read counts, and Sleuth-generated differential expression results are included as supplementary data (Tables S1- S3).

### Genomic mapping and metagene plots

High throughput sequencing reads were trimmed to remove adapters using bbduk.sh v39.13 with k=50, mink=10, hdist=1, minoverlap=8, minlength=20, trimq=10, tbo and tpe options. Then, trimmed reads were aligned to the GRCz11 zebrafish genome using STAR v2.7.10b with an Ensembl v113 GTF file. The --outWigType bedGraph option was used with STAR to allow visualization of coverage for each gene (Fig. S1A-B). To generate metagene plots, various modules from deepTools v3.5.6 were used. First, the bamCoverage module was used to generate strand specific counts per million normalized bigWig files containing the coverage at different genome positions for each sample. Then the computeMatrix module was used in scale-regions mode to produce intermediate files containing coverage for each gene scaled to a fixed length, as well as the coverage 1000 bases downstream of the genes. Two sets of regions/genes were defined at the computeMatrix step, one set containing genes that correspond with overexpressed transcripts in *MZpnrc2* mutants, and one set containing all other genes that were expressed in *MZpnrc2* mutants (i.e. non-overexpressed genes). The computeMatrixOperations utility was then used to combine the computeMatrix output for each strand into a single file. plotProfile was then run with --averageType mean as well as the – outFileNameData option in order to write the metagene coverage data to a file. In R v4.4.0, the coverage data was then averaged within each experimental condition, and the coverage was standardized so that the sum of the coverage for each condition/gene set combination summed to one across all positions on the metagene plot. The standardized coverage was then plotted using ggplot2 v3.5.2.

### uORF enrichment analysis

To identify transcripts with functional uORFs, we used a published dataset of zebrafish transcripts containing high-confidence uORFs from 24 hpf embryos (Johnstone et al., 2016). Among the time points sampled in Johnstone et al (2016), the 24 hpf dataset was the closest to mid-segmentation stages. Similar enrichment results were obtained using the 12 hpf dataset (Johnstone et al., 2016) (data not shown). To select for uORFs with evidence of translation, we followed methods in Gangras et al (2020) to filter the 24 hpf dataset (Johnstone et al., 2016) using the following criteria: 5′UTR RPF RPKM ≥ 5, RNA-Seq RPKM≥ 5, RPF RPKM ≥ 5, and ORF translation efficiency (RPF RPKM/RNA-Seq RPKM) >1. In addition to these criteria, the functional uORF list was filtered to only include APPRIS primary protein coding transcript isoforms that are functionally important (Rodriguez et al., 2022). Next, to identify functional uORF-containing transcripts in our dataset, we filtered our expression dataset (Table S3) to only include APPRIS primary protein coding transcript isoforms and compared this list with the filtered functional uORF list. A one-sided Fisher’s exact test was applied to evaluate the enrichment of transcripts with uORFs among the transcripts that were significantly overexpressed in *MZpnrc2* mutants (q < 0.05) versus transcripts that were non-overexpressed in *MZpnrc2* mutants (Table S3). hpf = hours post-fertilization; RPF = ribosome-protected mRNA fragment; RPKM = reads per kilobase mapped.

### Motif analysis

Motif analysis was performed using 3′UTR input sequences corresponding to transcripts identified from RNA-Seq analysis of *MZpnrc2* mutant embryos (Tables S3-S4) and the MEME Suite tool STREME version 5.5.6 (Bailey et al., 2021). Annotated 3′UTR sequences from each of the following datasets were analyzed using STREME: 1) significantly overexpressed transcripts (q < 0.05; n = 1320), 2) unaffected transcripts (log2FC from -0.05 to 0.05; n = 1334), and 3) significantly under-expressed transcripts (i.e. transcripts with decreased expression in *MZpnrc2* mutants relative to WT embryos; q < 0.05; n = 690) (Tables S3-S4). 3′UTR sequence annotations were obtained using BioMart (Ensembl release 113, GRCz11 assembly) and those less than 50 nts in length were removed to reduce annotation and length biases. Motif range was set to 6-8 nts and shuffled input sequences were used as the control with Markov order set to 2 (the default setting used by STREME). Sequence logos (Schneider and Stephens, 1990; Schneider et al., 1986) were generated using probability matrix data from STREME and the seqLogo package (Bembom and Ivanek, 2024) available with BioConductor version 3.19 (Gentleman et al., 2004; Huber et al., 2015) in RStudio version 2023.06.2 Build 561 (Posit Software). Motif enrichment results are reported in Table S5.

### GO analysis

GO analysis was performed using the PANTHER Overrepresentation Test (Released 2024-02- 26) that is available from the Gene Ontology Consortium at https://geneontology.org/ (Mi et al., 2013; Thomas et al., 2022). The GO Ontology database version used was DOI: 10.5281/zenodo.10536401 (Released 2024-01-17). The input list contains gene IDs for which at least one transcript was significantly overexpressed (q < 0.05) in *MZpnrc2* mutants and for which a GO annotation was identified (n = 1619 uniquely annotated GO IDs) (Table S4), while the reference list contains the full set of GO-annotated zebrafish genes in the database (n = 26,353). GO results are reported in Table S5. Statistical test type was a Fisher’s Exact test corrected by calculating the false discovery rate.

### In situ hybridization

Whole mount in situ hybridization was performed as previously described (Broadbent and Read, 1999; Jowett, 1998) using digoxygenin (DIG)-labeled riboprobes. Riboprobe for *her1* was previously described (Delaune et al., 2012; Dill and Amacher, 2005). Riboprobe for *rhov* was synthesized from a plasmid containing the *rhov* cDNA (cb832) received from the Zebrafish International Resource Center (ZIRC). *rhov*-containing plasmid was linearized with NotI-HF (NEB) and transcribed with T7 RNA polymerase (Roche Life Science) to make DIG-labeled antisense *rhov* riboprobe. Riboprobe templates for *sp5l*, *mespab*, *ripply2*, *dact1*, *spry1*, and *spry4* were generated by PCR amplification of coding sequence (primer sequences are listed in Table S6). All DIG-labeled PCR-based probes were synthesized by transcribing PCR templates with SP6 or T7 RNA polymerase (Roche Life Science). *dact1*, *sp5l*, and *spry4* produced expression patterns consistent with direct submission in situ hybridization data (Thisse et al., 2001; Thisse et al., 2005). Additionally, expression patterns for all PCR-based probes shown in Fig. 2 match published expression patterns for WT embryos at mid-segmentation stages, including *dact1* (Gillhouse et al., 2004), *mespab* (Cutty et al., 2012; Wanglar et al., 2014), *sp5l* (Superina et al., 2014), *ripply2* (Junker et al., 2014; Kinoshita et al., 2018; Moreno et al., 2008; Wanglar et al., 2014; Yabe et al., 2016), *spry1* (Komisarczuk et al., 2008), and *spry4* (Nguyen-Chi et al., 2012). In situ hybridized embryos were mounted in Permount and imaged using an Axiocam HRc digital camera with AxioPlan2 microscope and 10x objective (Zeiss).

### RNA extraction and cDNA synthesis

Whole embryos at mid-segmentation stages (n = 15 embryos per biological replicate for each genotype in Figs 2, S2, and S3; n = 10 embryos per biological replicate for each genotype and condition in Fig. 7) were solubilized in Trizol for RNA extraction and purified following standard procedures (Thermo Fisher). All genotypes and conditions were analyzed for a minimum of 3 biological replicates. 500 ng of total RNA was reverse transcribed with random primers (Figs 2 and S2) or oligo(dT) primers (Fig. S3) and Superscript IV reverse transcriptase according to the manufacturer’s instructions (Thermo Fisher).

### Quantitative RT-PCR analysis

Quantitative RT-PCR was performed using PowerUp SYBR Green Master Mix (Thermo Fisher) and 4.5 ul cDNA (diluted 1:50 in Figs 2 and S2, diluted 1:25 in Figs S3 and 7) in 20 ul reactions, following manufacturer’s procedures. Negative controls lacking template were included for each primer set. All reactions were subjected to thermal melting to confirm that each reaction gave single peaks. Transcript levels were normalized to *mobk13* (*mob4*) (Gangras et al., 2020; Hu et al., 2016; Tietz et al., 2020). Cycle thresholds (Ct) were determined using CFX Maestro version 2.3 software (Bio-Rad). The average Ct of 2 technical replicates per sample was used for all fold-calculations except for calculations in Fig 4 for which the average Ct of 3 technical replicates per sample was used to more accurately quantify RNA from low-abundance fractions.

Changes in mRNA expression were calculated by ΔΔCt = ΔCt target − ΔCt control. ΔCt values were calculated using the formula Ct target – Ct *mob4* for each sample except for calculations in Fig 4 for which the formula used was Ct target – Ct Luciferase (see methods for analysis of polysome fractions below). Relative changes in mRNA expression levels are represented graphically as fold change, where relative mRNA fold change = 2−ΔΔCt. All graphs and statistics were generated using Prism 10 (GraphPad).

### Poly(A) tail-length assay

RNA for poly(A) tail-length assays was obtained by collecting whole embryos at mid-segmentation stages (n = 18 embryos for each genotype) followed by solubilization in Trizol and purification following standard manufacturer procedures (Thermo Fisher). Two biological replicates were performed for each genotype and produced consistent results (data not shown). Tail-lengths for *her1, her7, dlc, rhov,* and *ddx61* transcripts were assessed using the Poly(A) Tail-Length Assay Kit following standard manufacturer procedures (Thermo Fisher). Minus RT controls were performed for all samples and no amplification was detected as expected (data not shown). Primers used for tail-length assay PCR are listed in Table S6. Gel images were captured using an AlphaImager system (ProteinSimple) with exposure times set just below saturation for each lane analyzed. Pixel intensity profiles were generated using the lane profile analysis tool in AlphaImager HP version 3.4.0 (build 0728) and exported as png files for manual tracing using the anchor point tool in Adobe Illustrator version 24.1.2. Peaks were identified using the auto-detection option that identifies the upper and lower thresholds for peaks. Lane profiles shown in Fig. 3B are scaled individually for each lane profile to better illustrate pixel intensity distributions, whereas lane profiles in Fig. S3A-E are universally scaled between WT (lane 2) and *MZpnrc2* (lane 3) profiles to better illustrate differences in pixel intensities between lanes. Molecular weights were calculated using a molecular weight tool in AlphaImager that utilizes known sizes of the molecular weight ladder to determine the size and range of PCR products of interest.

### Preparation and analysis of poly(A)-enriched and poly(A)-depleted extracts

RNA for comparison of expression levels in poly(A)-enriched versus poly(A)-depleted RNA fractions was obtained from the same RNA sources as was used for RNA-Seq experiments. Three biological replicates were obtained for both WT and *MZpnrc2* mutant embryos. From each sample, 1 ug of total RNA was adjusted to a final volume of 50 ul in bead binding buffer (20 mM Tri-HCl, pH 7.5, 1.0 M LiCl, and 2 mM EDTA) and incubated with Dynabeads Oligo(dT)_25_ following manufacturer procedures (Thermo Fisher). RNA bound to beads (poly(A)- enriched RNA) was eluted in 10 mM Tris-HCl (pH 7.5). Unbound RNA (poly(A)-depleted RNA) was precipitated by adding 0.1x volume 3M sodium acetate and 2x volume ice-cold ethanol, followed by vortexing and incubation overnight at -80 C. After centrifugation at 4 C for 20 minutes at 12,000 x g, precipitated pellets from unbound RNA fractions were rinsed with ice-cold 70% ethanol and resuspended in 10 mM Tris-HCl (pH 7.5) at an equivalent volume used for elution of bound RNA fractions. Each RNA fraction was reverse transcribed using random primers and Superscript IV reverse transcriptase (RT) according to the manufacturer’s instructions (Thermo Fisher). As a control, 1 ug of total RNA from the same samples used for bead-based fractionation were reverse transcribed in parallel. RT-qPCR analysis was performed as described earlier using 4.5 ul of cDNA diluted 1:10.

### Protein extraction and immunoblot analysis

Protein extracts were prepared from zebrafish embryos or larvae at 1-5 days post-fertilization. Individuals were collected and washed twice with rapid pipetting to disrupt yolks in 200 ul of ice-cold Ringer’s solution with protease inhibitors (116 mM NaCl, 2.9 mM KCl, 5.0 mM HEPES, pH 7.2, 100 μg/mL Soybean Trypsin Inhibitor [Sigma], 0.2 mg/mL PMSF, Complete Protease Inhibitor Cocktail [Roche]). Embryo and larval suspensions were centrifuged at 4 C for 1 minute at 300 x g for each wash. Cell lysis was performed by addition of 1 ul 4X LDS Buffer (Thermo Fisher) for each embryo or larva collected (n = 20-65 individuals per extraction), followed by homogenization with a plastic pestle, vortexing for 1 minute, and boiling for 10 minutes.

Homogenization, vortexing, and boiling were performed twice sequentially. Embryo or larval equivalents were loaded on a 4-12% Novex Bis-Tris polyacrylamide gel (Thermo Fisher), electrophoresed in NuPAGE MOPS SDS running buffer (Thermo Fisher) alongside a Novex Sharp pre-stained protein standard (Thermo Fisher), and transferred to a PVDF membrane using the Bio-Rad wet transfer system following standard manufacturer procedures. Immunoblot was blocked in 5% non-fat skim milk in Tris-buffered saline (20 mM Tris, 150 mM NaCl, pH 7.5) and subsequent steps were performed following standard procedures available online from Bio-Rad in bulletin 6376 (https://www.bio-rad.com/). The immunoblot was co-incubated with DDX6 (A16270, AbClonal) and β-actin (sc-47778, Santa Cruz Biotechnology) primary antibodies followed by co-incubation with AP-conjugated goat anti-rabbit IgG (H+L) secondary antibody (ThermoFisher) to detect DDX6 primary antibody and goat anti-mouse IgG (H+L) secondary antibody (Thermo Fisher) to detect β-actin primary antibody. In control experiments, immunoblot antibody incubations were performed separately and showed no differences in DDX6 and β-actin signal detection (data not shown). Primaries were diluted 1:1000 and secondaries were diluted 1:10,000 in blocking solution with 0.1% Tween-20. Primary antibody incubations were performed overnight at 4 C, secondary antibody incubations were performed for 1 hour at room temperature. Chemiluminescent detection was performed using CDP-Star following the manufacturer’s procedure (Thermo Fisher) and imaged using the Sapphire FL Biomolecular Imager (Azure Biosystems). Quantification of signal intensities was performed using lane profile analysis in AlphaImager HP version 3.4.0, build 0728 (Protein Simple).

### Polysome profiling

WT and *MZpnrc2* mutant embryos were raised to mid-segmentation stages and treated in 100 ug/ml cycloheximide for 5 minutes (n = 300 embryos per biological replicate for each genotype). Embryos were triturated using a 200 ul pipette and washed to remove yolks as previously described (Link et al., 2006) except that E2 embryo medium containing cycloheximide (15 mM NaCl, 0.7 mM NaHCO_3_, 0.5 mM KCl, 0.15 mM KH_2_PO_4_, 2.7 mM CaCl_2_, 0.5 mM Na_2_HPO_4_, 1mM MgSO_4_, 100 ug/ml cycloheximide) was used in lieu of 0.5x Ginzburg Fish Ringer’s buffer. Cell pellets were resuspended in 500 ul of chilled lysis buffer (20 mM Tris HCl pH 7.5, 30 mM MgCl_2_, 100 mM NaCl, 0.5 uM DTT, 100 ug/ml cycloheximide, 1 mg/ml heparin, 2.5 ul/ml NP-40) and incubated on ice for 10 minutes. Lysates were centrifuged at 12,000 x g for 10 minutes at 4°C to fractionate nuclei and debris (pellet) from cytoplasm (supernatant). After total RNA quantification of cytoplasmic extracts on a Nanodrop Spectrophotometer (Thermo Fisher), a volume corresponding to 50 ug of total RNA was transferred onto a 10-50% linear sucrose density gradient per sample. Gradients were centrifuged at 35,000 rpm in a SW41 rotor for 2-3 hours at 4°C. Following centrifugation, gradients were fractionated into 12, 1 ml fractions using a BioComp Piston Gradient Fractionator and A_260nm_ profiles were recorded using a BioComp Triax Flow Cell.

### RNA extraction and quantitative RT-PCR analysis of polysome fractions

From each fraction, 400 ul was used for the following procedure. To each fraction, 2 ng Luciferase Control RNA (L4561, Promega) was added as a normalization spike-in control, followed by 600 ul of Trizol and purification following standard procedures (Thermo Fisher). RNA pellets were rinsed with ice-cold 70% ethanol, briefly dried, and resuspended in 20 ul nuclease-free H_2_O. For each fraction, 8 ul of RNA was reverse transcribed using random primers and Superscript IV reverse transcriptase following the manufacturer’s instructions (Thermo Fisher). For quantitative PCR, 4.5 ul of cDNA (diluted 1:10) was used in 20 ul reactions with PowerUp SYBR Green Master Mix following the manufacturer’s instructions (Thermo Fisher). Transcript levels were normalized to Luciferase RNA to control for differences in RNA recovery across fractions. Luciferase (Fluc) primer sequences are listed in Table S6. Cycle thresholds (Ct) were determined using CFX Maestro version 2.3 software (Bio-Rad). The average Ct of 3 technical replicates per sample for each primer pair was used for all calculations. Relative proportions of mRNA abundance across fractionated samples (represented as percentages) were calculated following methods published by Pringle et al., (2018) for analysis of transcript abundance in fractioned RNA samples with normalization to Luciferase RNA. To compare differences in mRNA abundance in equivalent fractions from WT and *MZpnrc2* samples, relative differences in mRNA abundance in each fraction between wild-type and *MZpnrc2* mutant samples were calculated using the formula ΔΔCt = ΔCt target − ΔCt control, with the control being ΔCt values from WT extracts and the target being ΔCt values from *MZpnrc2* mutant extracts. ΔCt values were calculated using the formula Ct target – Ct Luciferase for each sample. Relative changes in mRNA expression levels are represented graphically as fold change, where relative mRNA fold change = 2−ΔΔCt. All graphs and statistics were generated using Prism versions 8 and 10 (GraphPad).

### Ddx6 immunolabelling and imaging

Embryos at mid-segmentation stages were fixed in 4% PFA in PBS overnight at 4 C, and washed 5 x 5 minutes in PBS with 0.1% Tween-20 (PBST). Embryos were then permeabilized in 1X PBS containing 2% Triton-X for 2 hours at room temperature, followed by 3 x 5 minutes PBST washes. Blocking was performed in PBST with 2% BSA, 2% goat serum for 1 hour at room temperature, followed by incubation with 1:200 dilution anti-DDX6 (A16270, AbClonal) in PBS with 2% BSA, 2% goat serum, 1% DMSO and 0.1% Tween-20 (PBST) overnight at 4 C. Embryos were washed 8 x 15 minutes in PBST, blocked as described above, followed by incubation with 1:400 dilution Alexa Fluor 488 goat anti-rabbit secondary antibody (Thermo Fisher) in PBS with 2% BSA, 2% goat serum, 1% DMSO and 0.1% Tween-20 (PBST) overnight at 4 C. Embryos were washed 8 x 15 minutes in PBST and stored in SlowFade Gold Anti-fade Mountant with DAPI (S36939, Thermo) prior to imaging. Immunolabelled embryos were deyolked, dissected, and mounted in Fluoromount-G and imaged with a Plan VC 60X objective on an Andor Revolution WD spinning disk confocal system with Nikon TiE inverted microscope using MetaMorph software (version 7.10.4.407) and a pco.edge 4.2 bi 16-bit sCMOS camera (Excelitas). 488 and 405 laser intensities were set to 20% and exposure time set to 2000 ms per Z-stack slice per channel. For each specimen, the range of Z-stack slices that best captured the presomitic mesoderm (PSM) was determined by setting the upper and lower bounds of the Z-stack to match the upper and lower bounds of the most proximal, fully formed somite that lies in the same plane as the PSM when viewed laterally (See Fig. 5C).

### Confocal imaging analysis

Images were analyzed using the package Fiji (ImageJ version 1.54p) (Schindelin et al., 2012; Schneider et al., 2012). Three z-stack slices, separated by 2 um intervals per slice, were analyzed for each sample to comprehensively analyze Ddx6/Ddx61 immunolabelling in the presomitic mesoderm (PSM). Of the three slices, quantification of central slices is shown in Fig. 5 and quantification of higher and lower z-stack slices is shown in Fig. S4. For mean intensity and puncta quantifications, the brightest image across all images being compared was used to set the maximum signal intensity across all samples analyzed to avoid saturated, overexposed pixels. A single region of interest (ROI) that best captures the PSM of a single specimen was manually drawn and kept constant across all images analyzed and re-positioned when necessary to ensure that all quantifications were performed on identically sized ROIs that best captured the PSM. The mean intensity of each ROI was calculated using the measure analysis tool in Fiji. To quantify puncta number within ROIs, the “find maxima” tool was used with prominence set to 5000 which best captured the range of puncta sizes across all samples relative to lower or higher prominence settings that over- or under-counted visible puncta, respectively (data not shown).

### Plasmid construction & transgenesis

The heat shock construct *hsp70l:Myc-cnot7DN-SV40 pA* was assembled by first constructing plasmid *pMA-Myc-cnot7DN* using the GeneArt Plasmid Construction Service (Thermo Fisher). *pMA-Myc-cnot7DN* contains the coding sequence for a dominant negative form of zebrafish *Cnot7* (*Cnot7DN*) with an N-terminal C-myc epitope tag (Evan et al., 1985) and intervening SGLRS flexible linker peptide. The *myc*-tagged *Cnot7DN* coding sequence was based on plasmid M338 (*pCS2 + MT-CNOT7-D40A,E42E,C67E,L71E-sv40*) that has previously been used to inhibit deadenylase function in zebrafish embryos (Fujino et al., 2018; Makino et al., 2015; Mishima and Tomari, 2016; Mishima and Tomari, 2017). After sequence validation of *pMA-Myc-cnot7DN*, the *Myc-cnot7DN* insert was cloned into *pBSKI2* (Thermes et al., 2002) using standard restriction digestion-based cloning with the enzymes NheI-HF and BamHI-HF (NEB). Plasmid and I-SceI enzyme were injected into the blastomere of 1-cell stage WT and *MZpnrc2* mutant embryos following standard transgenesis procedures (Thermes et al., 2002). To determine a suitable dose, we injected 20 pg and 60 pg plasmid doses in the absence of heat-shock induction and assessed morphology at mid-segmentation stages; the 20 pg dose was well-tolerated (73 of 90 injected embryos had WT morphology), but the 60 pg dose caused gross abnormalities (0 of 56 injected embryos had WT morphology).

### mRNA injection

The *Myc-cnot7DN* coding sequence from plasmid *pMA-Myc-cnot7DN* was PCR amplified and cloned into expression vector pCS2+ (Rupp et al., 1994; Turner and Weintraub, 1994) using standard digestion-based cloning with the enzymes EcoRI-HF and XbaI (NEB) to generate the plasmid *pCS2-Myc-cnot7DN*. For deadenylation inhibition experiments, *Myc-cnot7DN* mRNA was synthesized using the SP6 mMessage Machine Kit (Thermo Fisher), diluted in 0.2M KCl with 0.1% phenol red, and injected into 1-cell stage embryos at either a 200 pg dose or a 400 pg dose of mRNA per embryo. Primer sequences are listed in Table S6.

### Heat shock assay

Plasmid-injected embryos (were collected and raised to segmentation stages (10-12S), heat shocked at 37 C for 30 minutes, and monitored for somite defects at 30 minute intervals post-heat shock (pHS). Because *MZpnrc2* mutants are developmentally delayed relative to WT embryos by ∼2-3 hours, heat shock induction was performed on stage-matched embryos from WT and *MZpnrc2* clutches based on somite number, requiring staggered heat shocks beginning with WT embryos followed by heat shocks 2-3 hours later for *MZpnrc2* mutant embryos when they reached the same somite stage. Live embryos at 6 hours post-heat shock were mounted in 3% methylcellulose and imaged at 63x magnification using AxioVision Software (Zeiss) on a MZFLIII Fluorescence Stereo Microscope (Leica) with an AxioCam HRc digital camera (Zeiss) and subsequently fixed in 4% PFA for anti-Myc immunohistochemistry.

### Anti-Myc immunohistochemistry

Embryos injected with *hsp70l:Myc-cnot7DN* plasmid that were heat shocked at ∼14 hours post fertilization (hpf) for 30 minutes were collected 6 hours post-heat shock (∼22 hpf) alongside non-heat shocked controls, fixed in 4% PFA in PBS overnight at 4 C, and washed 5 x 5 minutes in PBS with 0.1% Tween-20 (PBST). Embryos were then incubated in water for 5 minutes, permeabilized with ice-cold acetone for 7 minutes, and incubated again in water for 5 minutes followed by 3 x 5 minute PBST washes. Blocking was performed in PBST with 2% BSA, 2% goat serum for 1 hour at room temperature, followed by incubation with 1:500 dilution monoclonal anti-myc (clone 9E10, MA1-980, Thermo Fisher) in PBS with 2% BSA, 2% goat serum, 1% DMSO and 0.1% Triton X-100 (PBDTx) overnight at 4 C. Embryos were washed 8 x 15 minutes in PBST, blocked as described above, followed by incubation with 1:200 dilution Alexa Fluor 546 goat anti-mouse secondary antibody (Thermo Fisher) in PBDTx overnight at 4 C. Embryos were washed 8 x 15 minutes in PBST prior to imaging. Immuno-stained embryos were mounted in Fluoromount-G and imaged at 63x magnification using AxioVision Software (Zeiss) on an MZFLIII Fluorescence Stereo Microscope (Leica) with a mCherry filter set and AxioCam HRc digital camera (Zeiss).

### CRISPR-Cas9 mutagenesis of *ddx6* and *ddx61*

CRISPR RNA (crRNA) guide sequences were designed using Integrated DNA Technologies (IDT) custom design tool (https://www.idtdna.com/site/order/designtool/index/CRISPR_CUSTOM). crRNA sequences with the highest on-target scores with no predicted off-target sites were chosen. The impact of targeted mutations on gene function was maximized by targeting regions within exons encoding the DEAD box helicase domain (see Fig. S5A-B). Reagents for CRISPR-Cas9 mutagenesis were prepared as previously described for efficient mutagenesis (Klatt Shaw & Mokalled, 2021). For double knockdown experiments, crRNAs targeting *ddx6* and *ddx61* (25 uM each, 50 uM total) and trans-activating CRISPR RNA (tracrRNA) (IDT) (50 uM) were combined equally and annealed to generate dual guide RNA duplexes (dgRNA), followed by 1:1 dilution in duplex buffer (IDT) for a final concentration of 25 uM dgRNA. Annealed dgRNAs were next combined equally with 25 uM Cas9-NLS (QB3 MacroLab, UC Berkeley) that had been diluted from a 40 uM stock solution using Cas9 dilution buffer B (IDT). Final injection solutions contained 0.1% phenol red dye, 0.2 M potassium chloride, and 5 uM each of dgRNA and Cas9-NLS. Injection solutions were incubated for 5 minutes at 37 C to facilitate ribonucleoprotein complex (RNP) formation. AB wild-type embryos were injected at the 1-cell stage with 2 nl of injection solution. crRNA sequences are listed in Table S6. For single knockdown experiments, the same method described for double knockdown experiments was employed except that twice the sgRNA dose as double knockdown experiments was used to ensure that the total molar amount of gRNA injected was equivalent between experiments (i.e 2.5 uM each of *ddx61* + *ddx6* gRNAs or 5 uM *ddx61* sgRNA or 5 uM *ddx6* gRNA). Live embryos were mounted in 3% methylcellulose and imaged at 40X-63x magnification using AxioVision Software (Zeiss) on a MZFLIII Fluorescence Stereo Microscope (Leica) with an AxioCam HRc digital camera (Zeiss)

## Acknowledgements

We thank the Ohio State Zebrafish Facilities staff for excellent care, the Ohio Supercomputer Center for computational resources, the Genomics Shared Resource at the Ohio State Comprehensive Cancer Center for RNA analysis and DNA sequencing, Paula Monsma and the Ohio State Neuroscience West Campus Imaging Core, and the Ohio State Center for RNA Biology for support and resources. We thank Susan Cole, Kenneth Poss, Mira Pronobis, and Guramrit Singh for advice and critical comments. We thank Anton Blatnik, Geremy Lerma, Lauren Levesque, Benjamin Pastore, and Zhongxia Yi for advice.

## Data and resource availability

RNA-Seq datasets and additional data files are available at https://www.ncbi.nlm.nih.gov/geo under accession GSE319255. All additional data and resources are included in the article and supplementary information.

## Author contributions

T.L.G., M.C.B., C.C.A, K.G.T, D.M.P, A.M., and S.L.A. performed experiments. T.L.G., M.C.B., R.D., M.G.K., and S.L.A. analyzed data. All authors contributed intellectually and discussed the data and manuscript. T.L.G. wrote the manuscript and co-authors participated in editing.

