## Supplementary Materials - Figures, legends, methods for "The RNA binding proteins Ddx6 and Ddx61 support development in mRNA-decay deficient *pnrc2* mutants"

Fig. S1 (goes with Fig. 1)

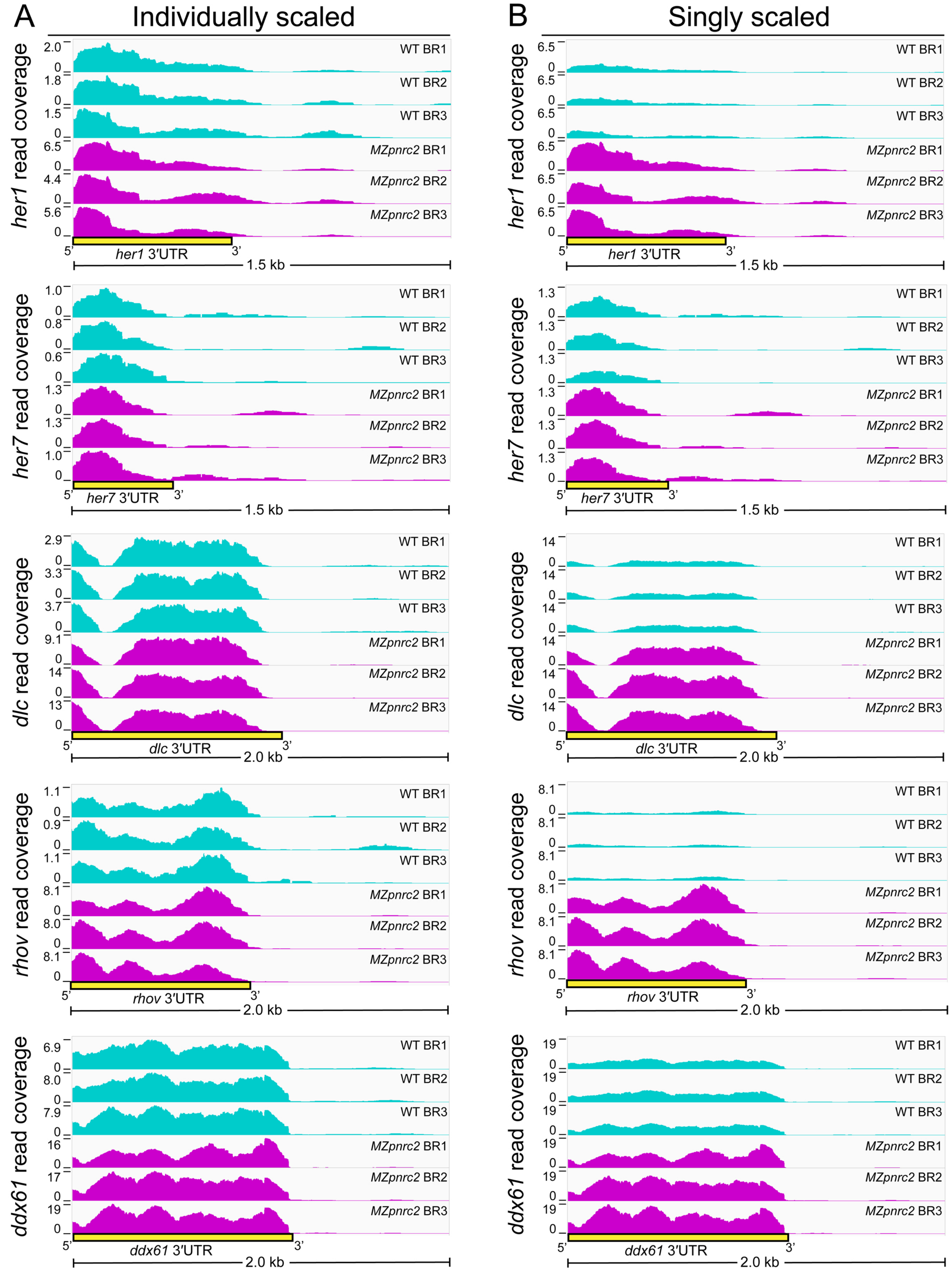

Fig. S1 (goes with Fig. 1)

C

#### Overexpressed - Biological Process

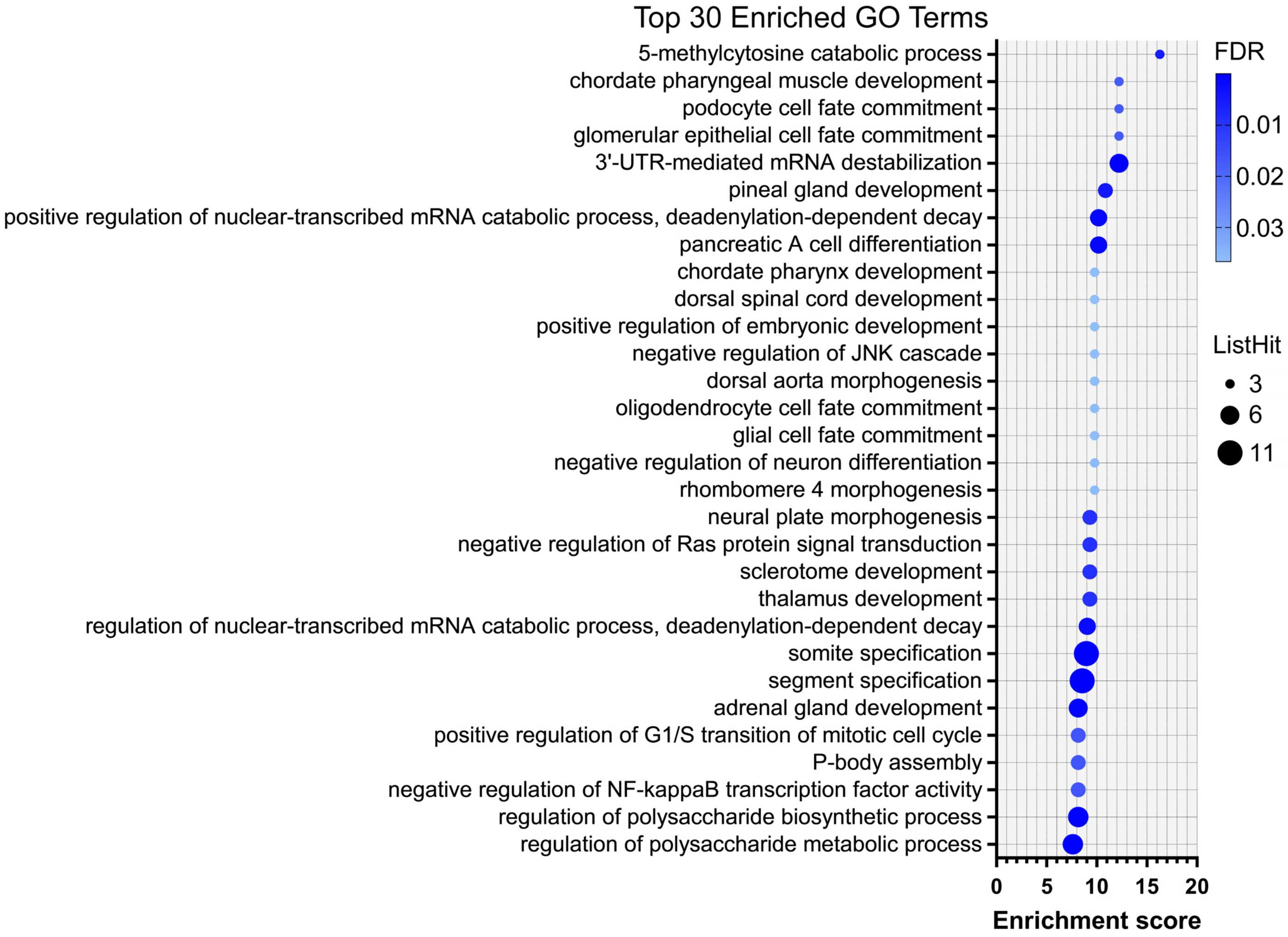

Fig. S1 (goes with Fig. 1)

D

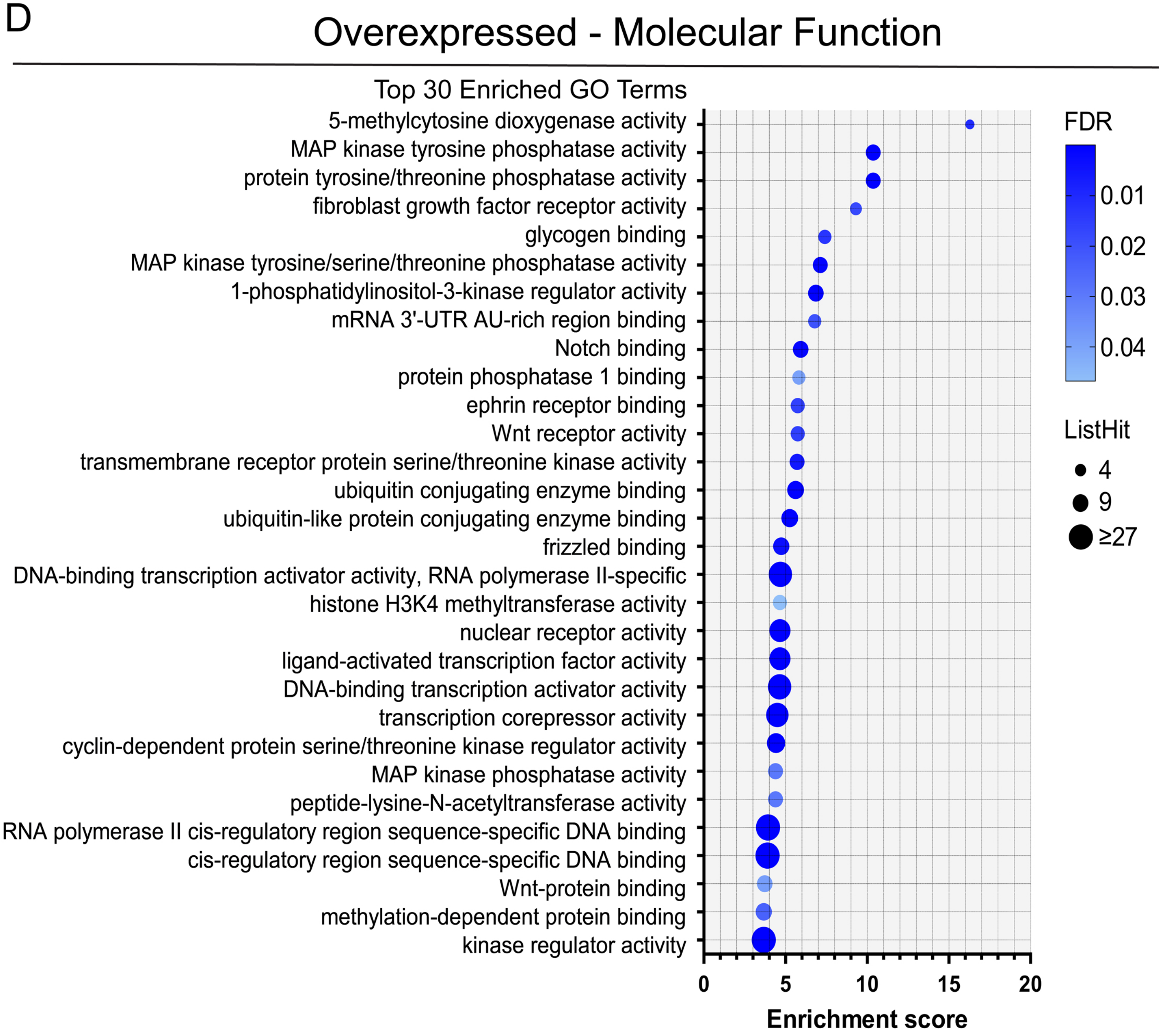

Fig. S1 (goes with Fig. 1)

E

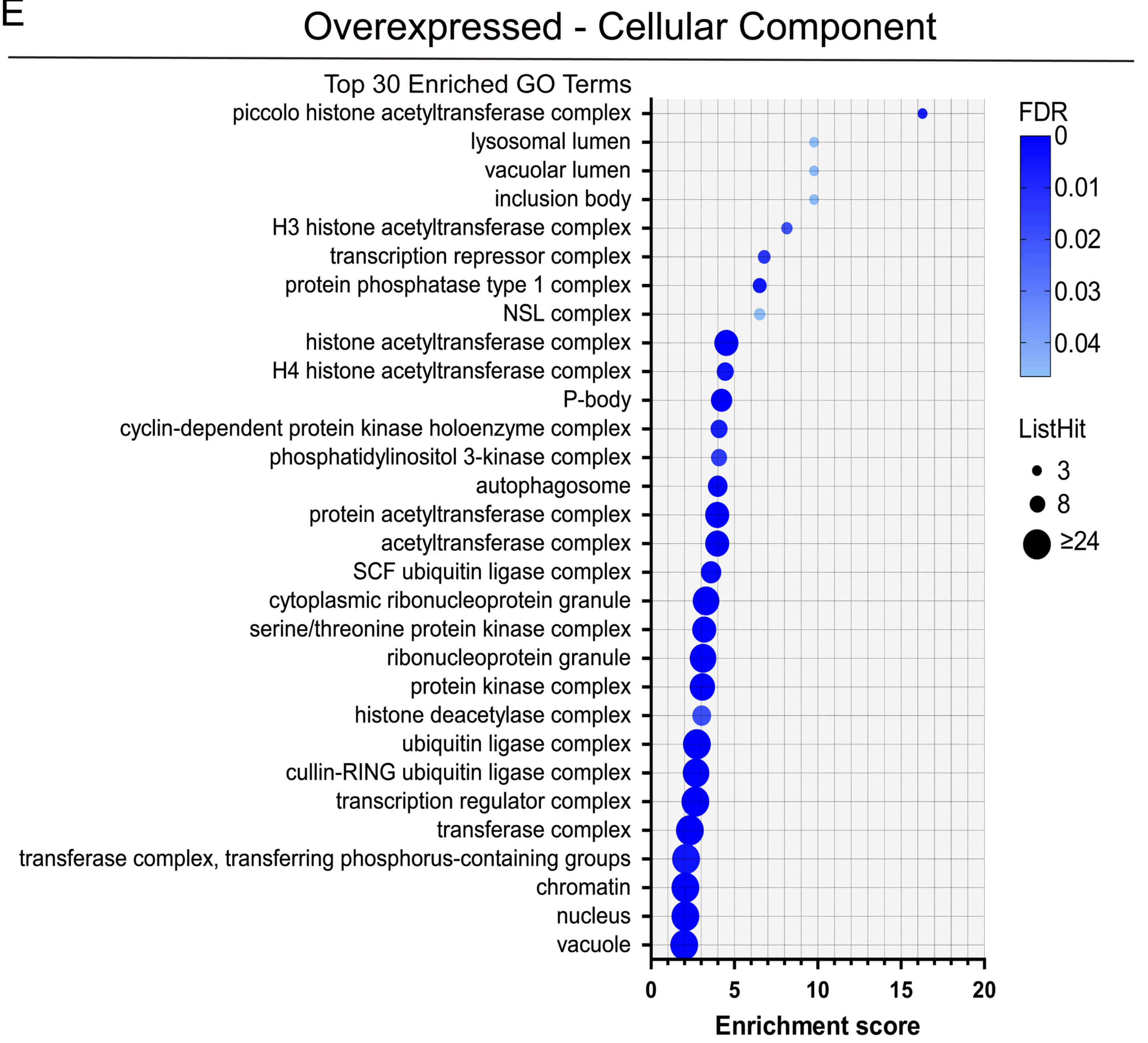

Fig. S1 (goes with Fig. 1)

F

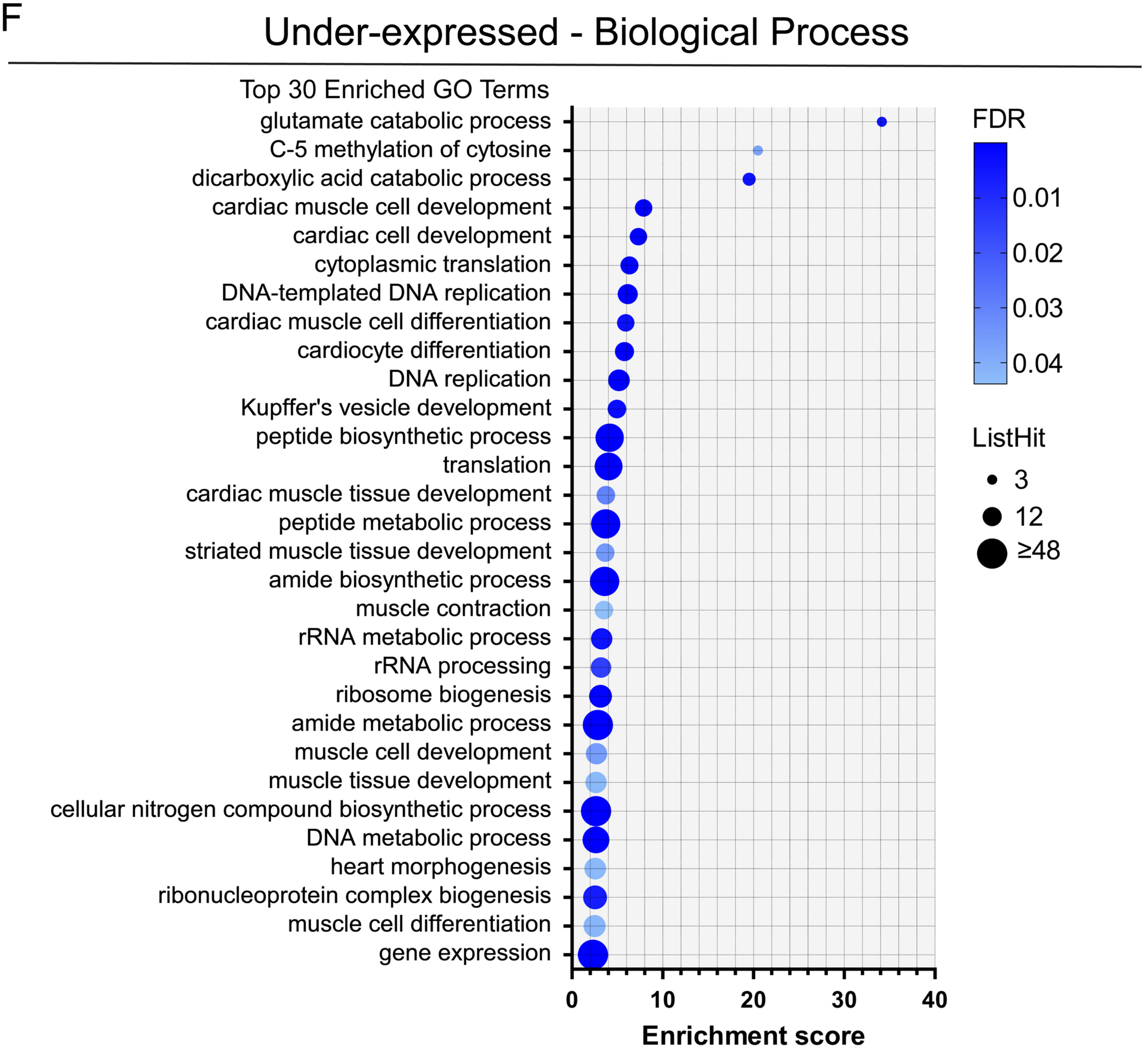

Fig. S1 (goes with Fig. 1)

G

#### Under-expressed - Molecular Function

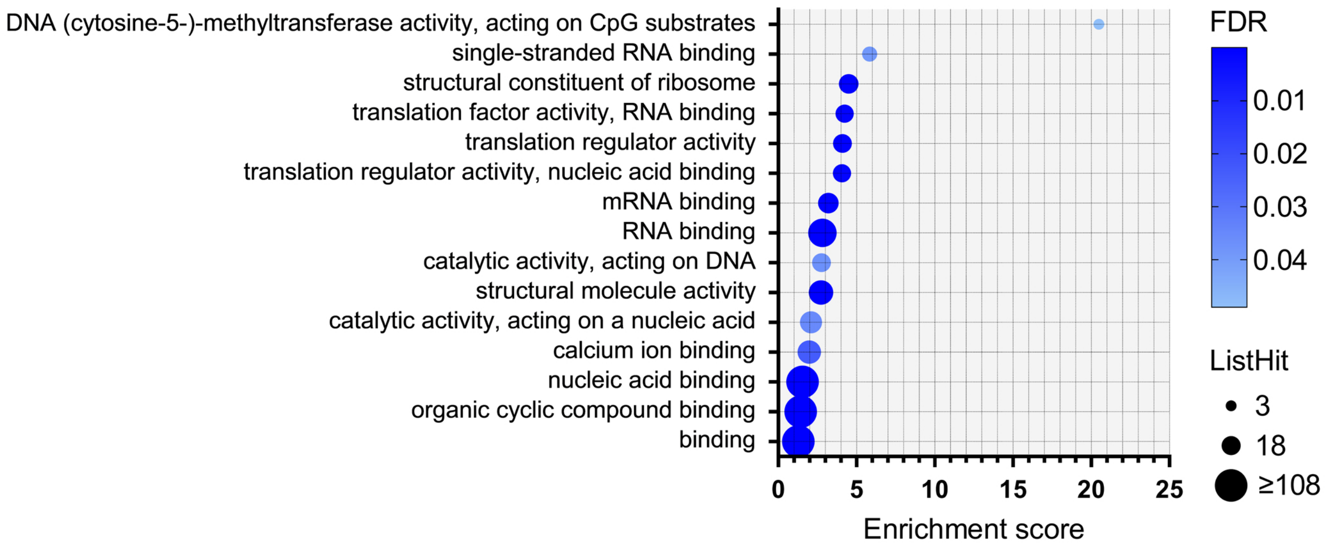


H

#### Under-expressed - Cellular Component

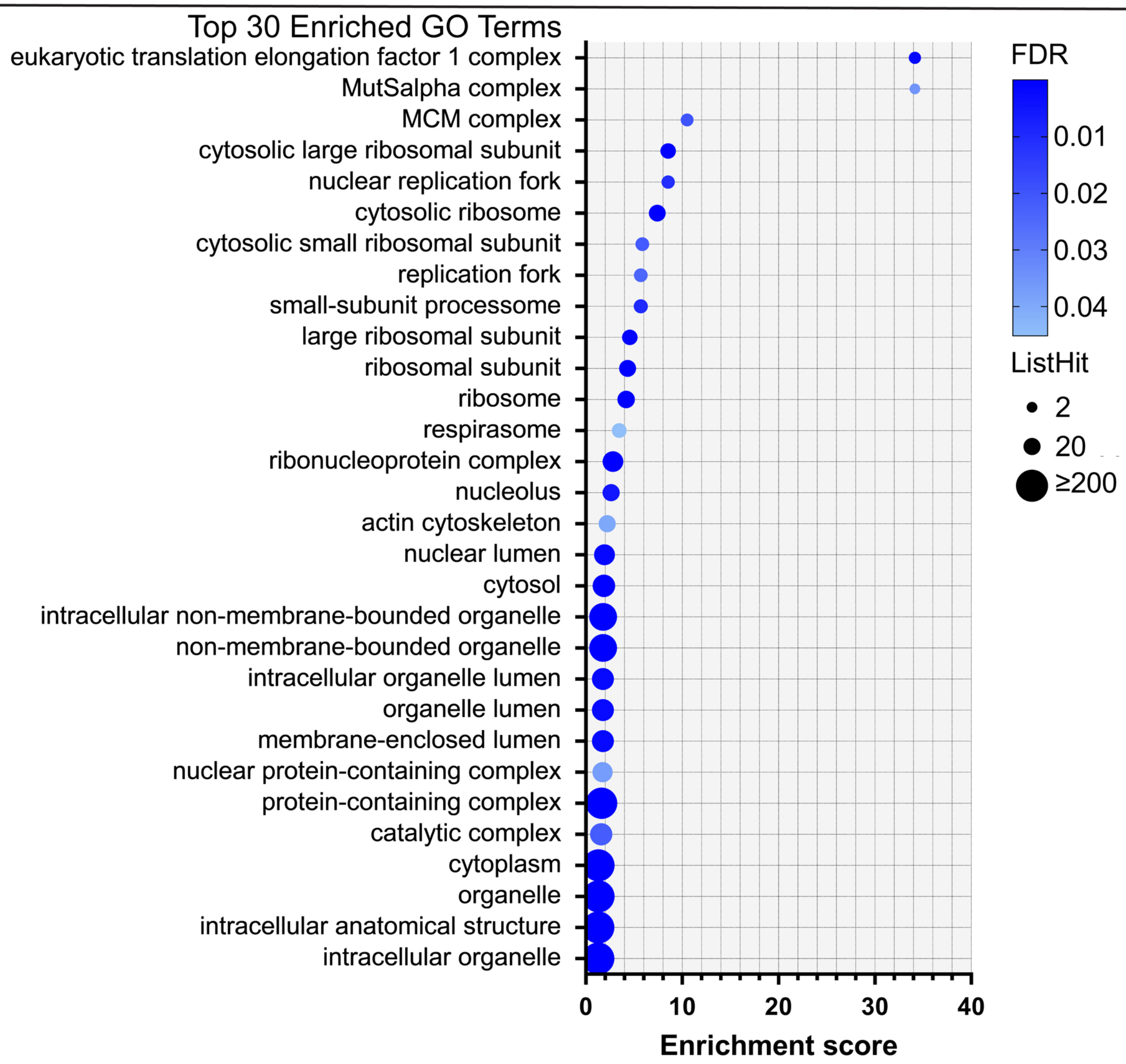

#### Supplementary Figure Legends

Fig. S1 (goes with Fig. 1). Read coverage analysis reveals that oscillatory gene transcript 3'UTR lengths are unaffected in *MZpnrc2* mutants and GO analysis of overexpressed genes in *MZpnrc2* mutants reveals many enriched GO terms. (A-B) Oscillatory gene transcript 3′UTR lengths are similar in WT and *MZpnrc2* mutants. Normalized bedgraphs (read coverage density plots) from high throughput sequencing reads (see methods in main text) of WT and *MZpnrc2* mutants at mid-segmentation stages (n = 50 embryos for each genotype per biological replicate). Read coverage is individually scaled in A to better show read density distribution across 3′UTRs and downstream regions for each bio-replicate, whereas read coverage is singly scaled across bio-replicates in B to better show differences in read density that correspond with transcript accumulation in *MZpnrc2* mutants. (C-E) Overexpressed transcripts in *MZpnrc2* mutants at mid-segmentation stages are enriched for GO terms associated with developmental pathways and post-transcriptional mRNA regulation. Top 30 most-enriched terms within the aspects biological process (C), molecular function (D), and cellular component (E) using GO enrichment analysis of overexpressed genes in *MZpnrc2* mutant embryos. (F-H) Under-expressed transcripts in *MZpnrc2* mutants at mid-segmentation stages are enriched for GO terms associated with translation. Top 30 most-enriched terms within the aspect biological process (F), all enriched terms within the aspect molecular function (G), and top 30 most-enriched terms within the aspect cellular component (H) using GO enrichment analysis of under-expressed genes in *MZpnrc2* mutant embryos. FDR = false discovery rate; ListHit = number of overexpressed genes per GO term.

Fig. S2 (goes with Fig. 2)

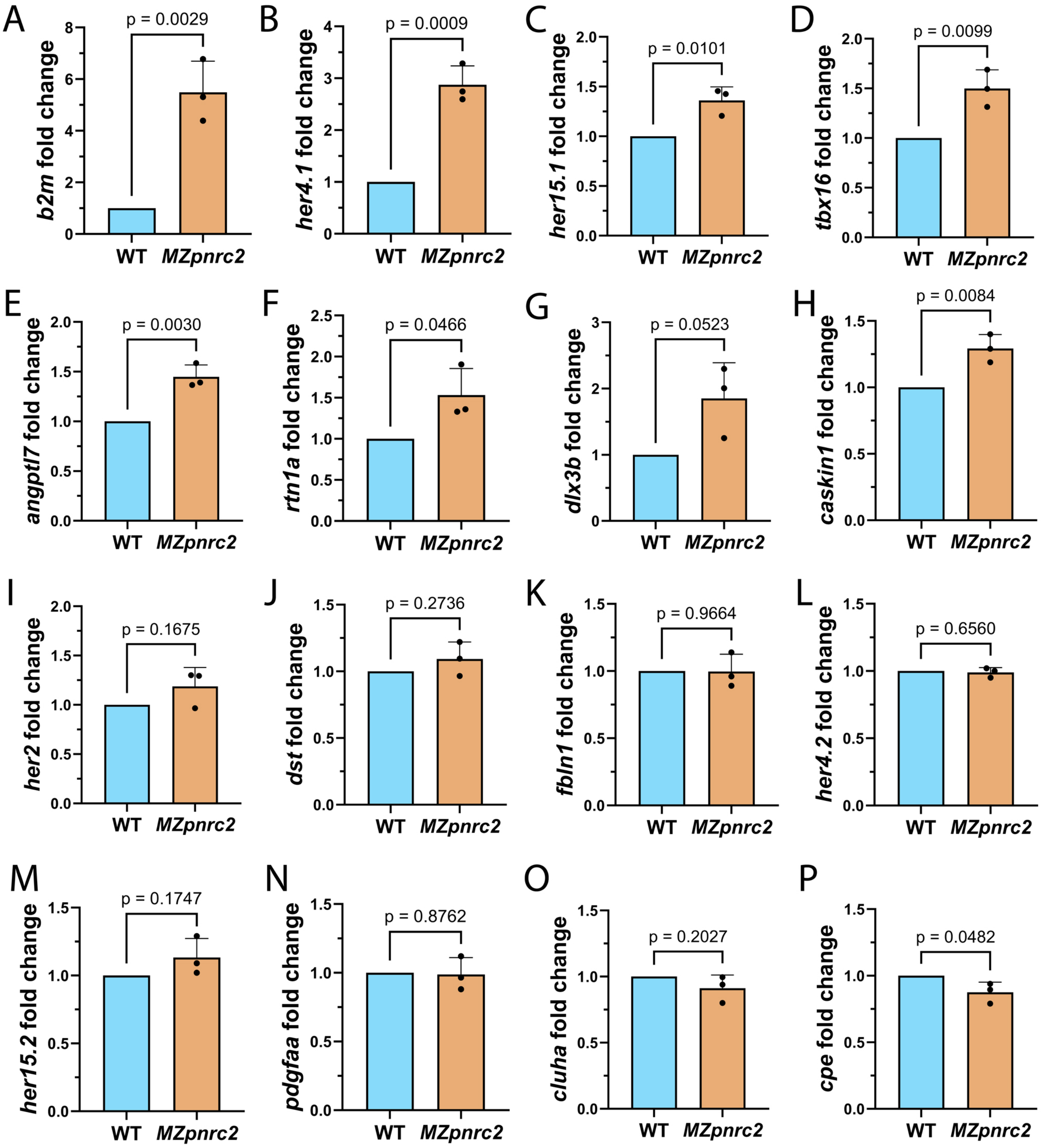

Supplementary Figure Legends

Fig. S2 (goes with Fig. 2). Many, but not all, genes known to oscillate in the zebrafish presomitic mesoderm are Pnrc2-regulated. (A-P) Bar graphs showing RT-qPCR validation of RNA-Seq results. Quantification was performed using cDNA synthesized from total RNA extracts from WT and *MZpnrc2* mutants at mid-segmentation stages (n = 15 embryos per biological replicate per genotype with 2-3 technical replicates per biological replicate). Fold change values are the means of biological replicate values derived from the average of technical replicates. P-values were calculated using an unpaired t-test. Among the overexpressed oscillatory gene transcripts shown, 4 were overexpressed in RNA-Seq analysis with false discovery rates of $q < 0.05$ (A-D), whereas 2 validated transcripts had larger q-values: *angptl7* ($q = 0.12$) (E) and *rtn1a* ($q = 0.13$) (F). The oscillatory gene transcript *dlx3b* was overexpressed in *MZpnrc2* mutants, but it did not reach significance in RNA-Seq ($q = 0.17$) nor RT-qPCR analyses (G). An additional transcript, *caskin1*, was overexpressed by RT-qPCR, but not in RNA-Seq analysis (FC = 0.96, $q = 0.18$) (H). One transcript, *her2*, was overexpressed in RNA-Seq analysis, but not by RT-qPCR (I). An additional 7 oscillatory gene transcripts were not overexpressed in RNA-Seq analysis nor by RT-qPCR analysis (J-O), whereas one transcript, *cpe*, was under-expressed in RNA-Seq analysis (FC = 0.83, $q = 0.08$) and validated as being under-expressed by RT-qPCR (P).

### Fig. S3 (goes with Fig. 3) - page 1

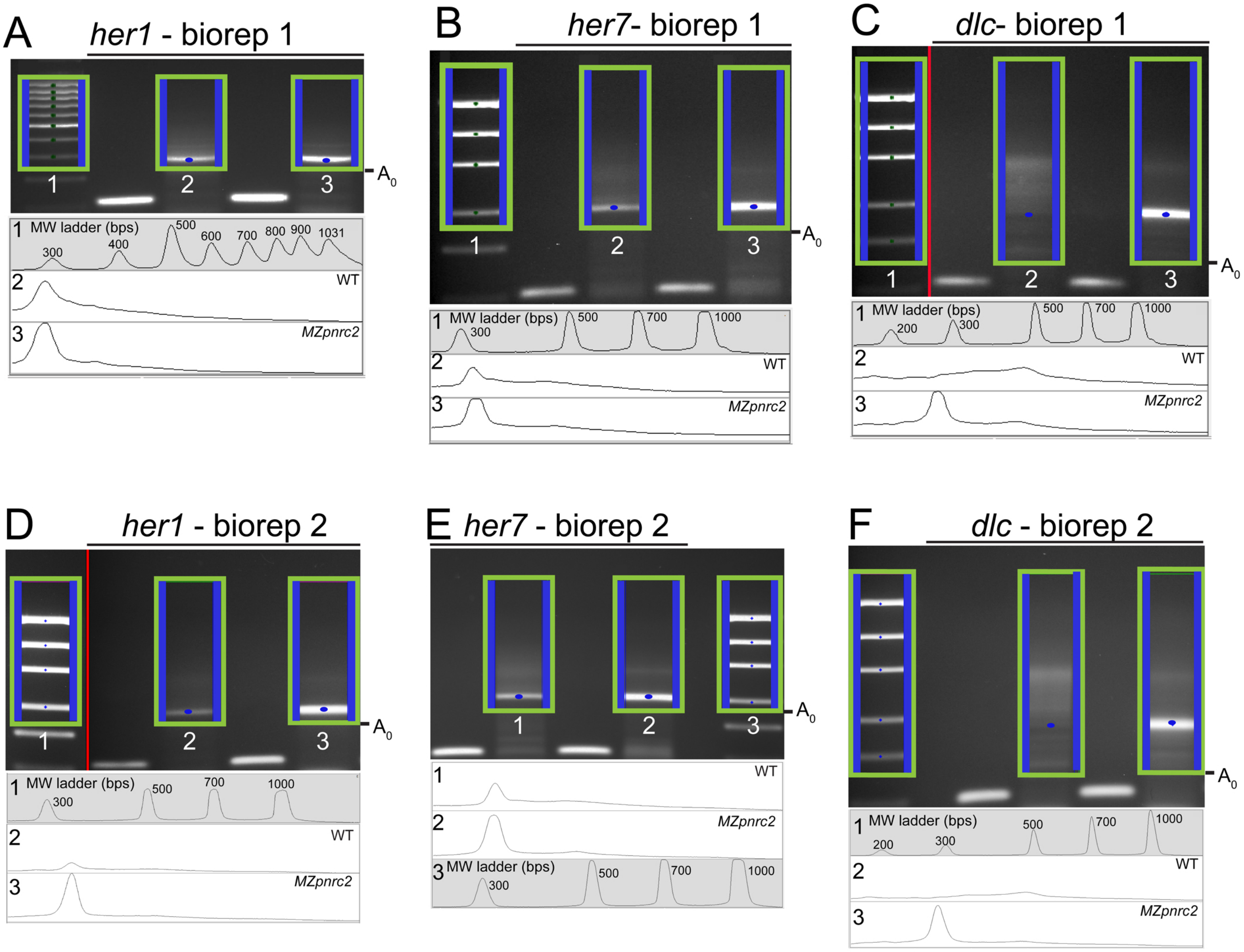

### Fig. S3 (goes with Fig. 3) - page 2

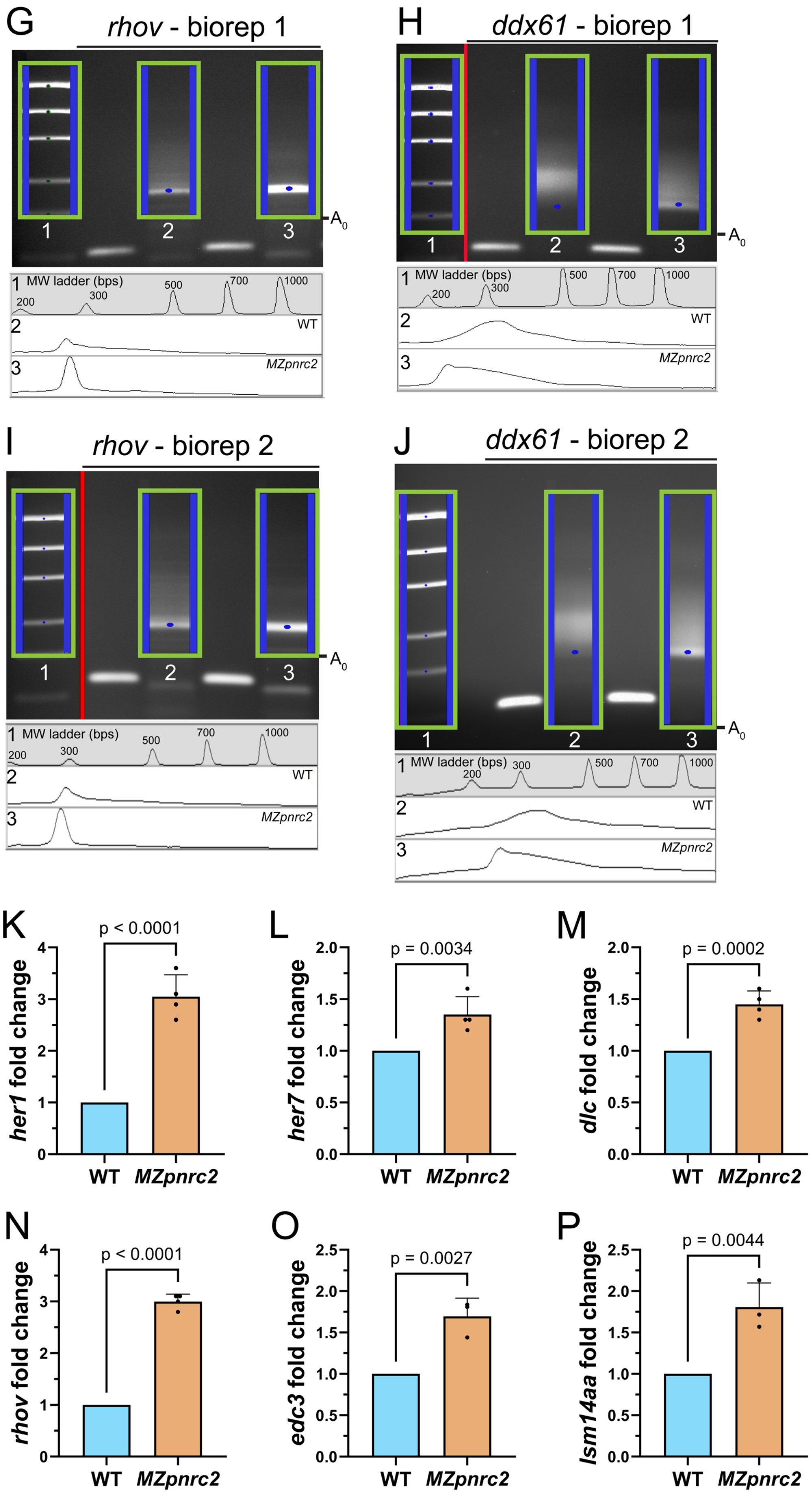

Supplementary Figure Legends

Fig. S3 (goes with Fig. 3). Overexpressed oscillatory gene transcripts and *ddx61* transcripts have shortened tail lengths in *MZpnrc2* mutants, but they are not completely deadenylated and coincide with transcript overexpression of additional P-body-associated factors. (A-J) Accumulated oscillatory gene transcripts as well as *ddx61* transcripts have shortened poly(A) tails in *MZpnrc2* mutants, but are not completely deadenylated. Larger images of gels shown in Fig. 3 are shown in A-C and G-H, with a second biological replicate shown for each poly(A)-tail length assay in D-F and I-J. Molecular weight ladders are shown for all tail length assays (note that for gel images that have been cropped and stitched so that the ladder is adjacent to profiled lanes for better size determination, stitched regions are denoted with a red line (C-D, H and I). Regions of interest for pixel intensity profiling are boxed in green, with blue lines denoting the width of each lane analyzed in pixel intensity plots shown below each gel. Predominant peaks are denoted with blue dots on each gel. Because *dlc* and *ddx61* lack a predominant peak in WT samples, a region corresponding to the peak in *MZpnrc2* mutants was identified by molecular weight and denoted with a blue dot in WT lanes. Gel images were captured under exposure times that were not saturated. Lane profiles of pixel intensities are scaled universally between lanes to better show differences in levels between WT and *MZpnrc2* mutants, although it is important to note that pA-tail reaction products are derived from end-point PCR where precise quantification is not possible. (K-P) Bar graphs showing quantification of oscillatory gene transcripts (K-N) and transcripts encoding P-body-associated factors (O-P) using cDNA synthesized with oligo(dT) primers (K-N) or with random primers (O-P) and total RNA extracts from WT and *MZpnrc2* mutants at mid-segmentation stages (n = 15 embryos per biological replicate, 2 technical replicates per biological replicate). Fold change values are the means of biological replicate values derived from the average of two technical replicates. P-values were calculated using an unpaired t-test.

### Fig. S4 (for Fig. 5) - page 1

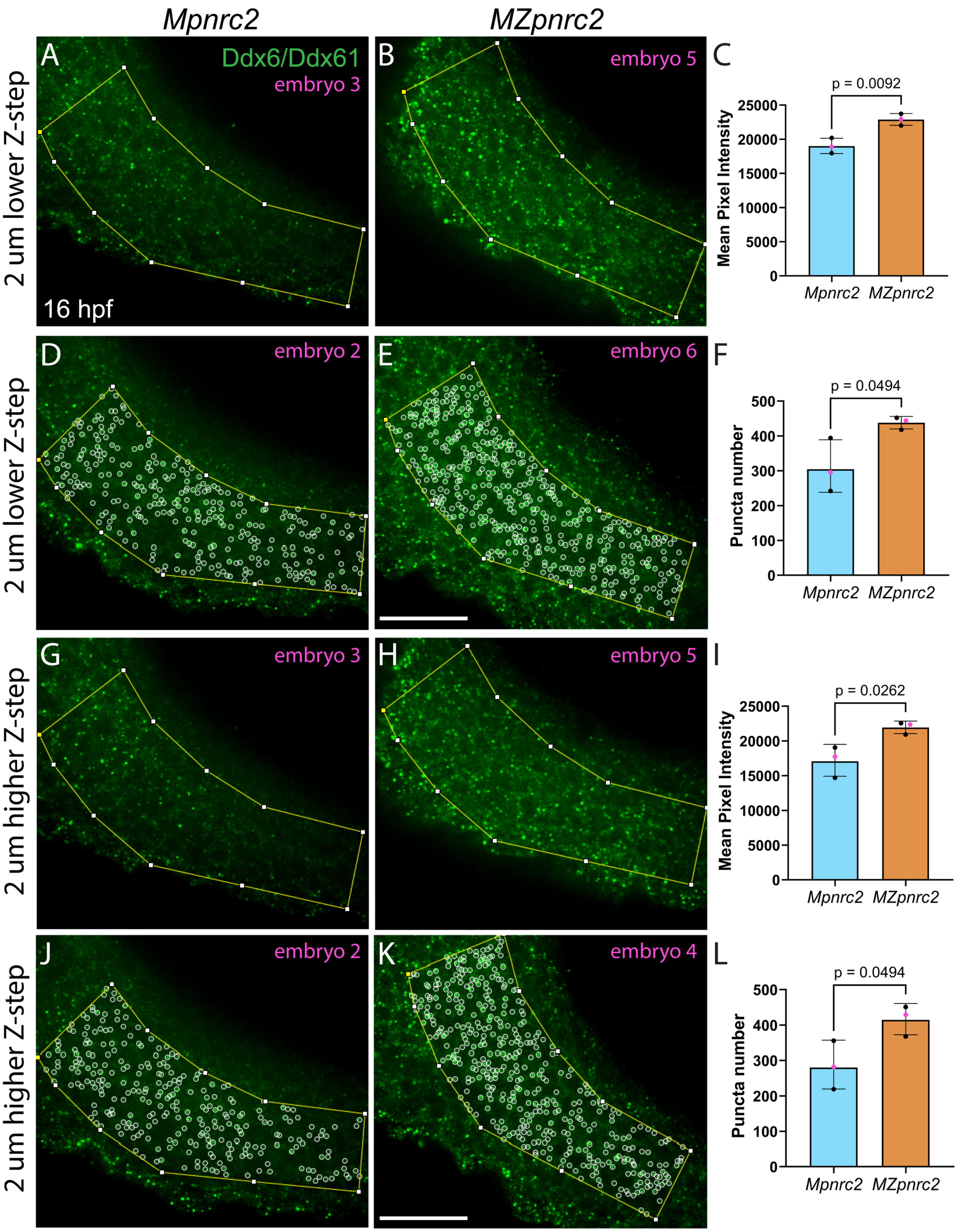

Fig. S4 (for Fig. 5) - page 2

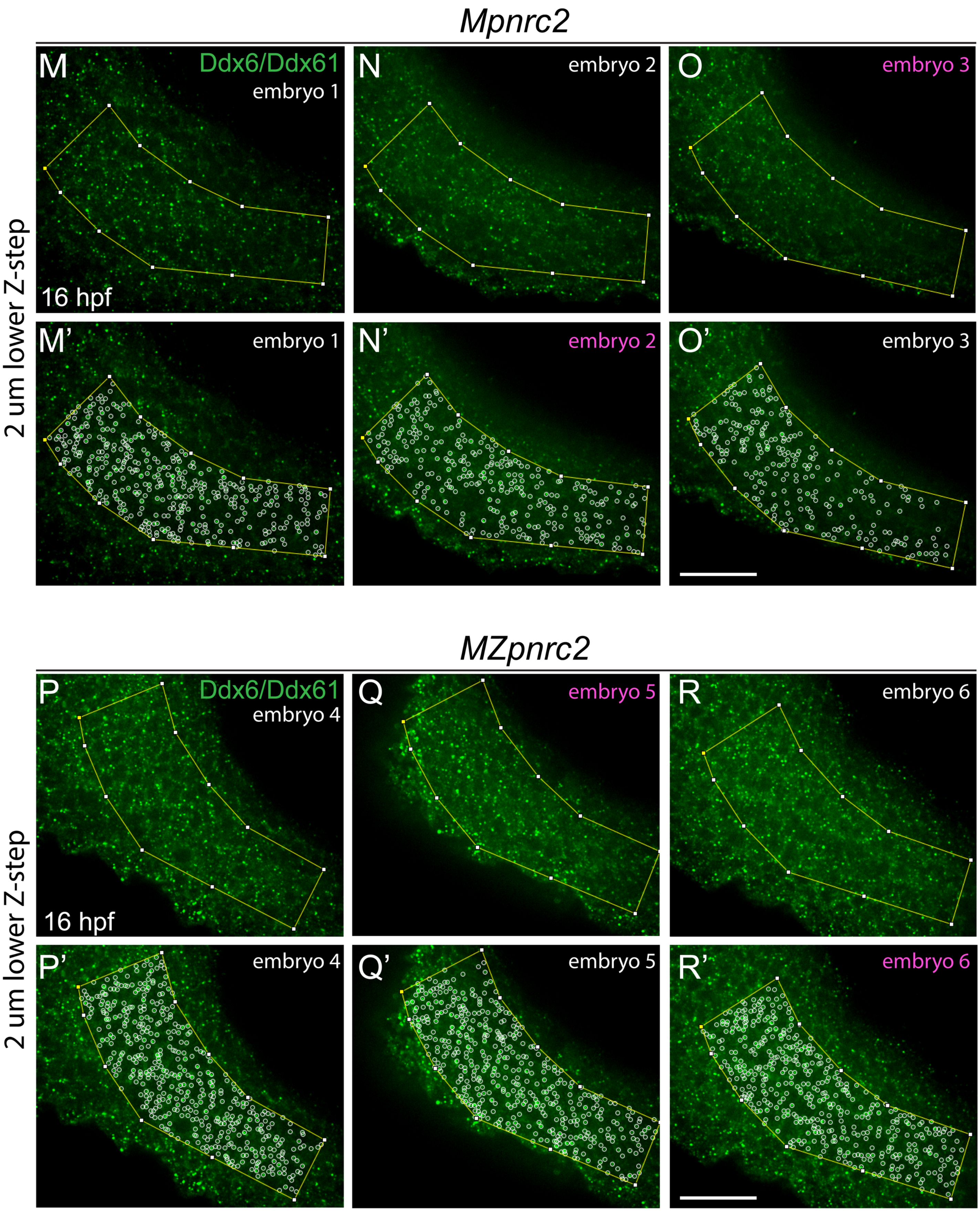

Fig. S4 (for Fig. 5) - page 3

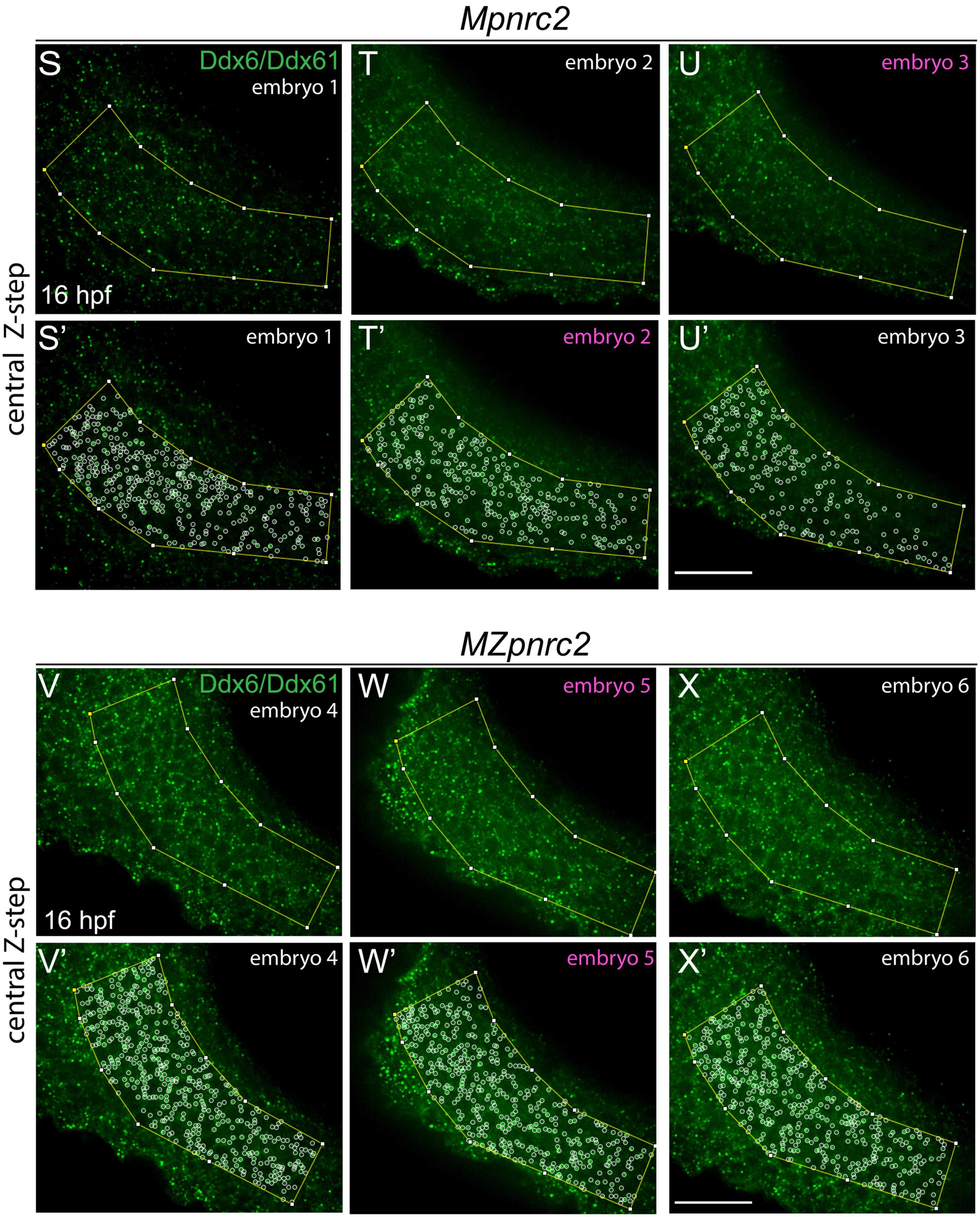

Fig. S4 (for Fig. 5) - page 4

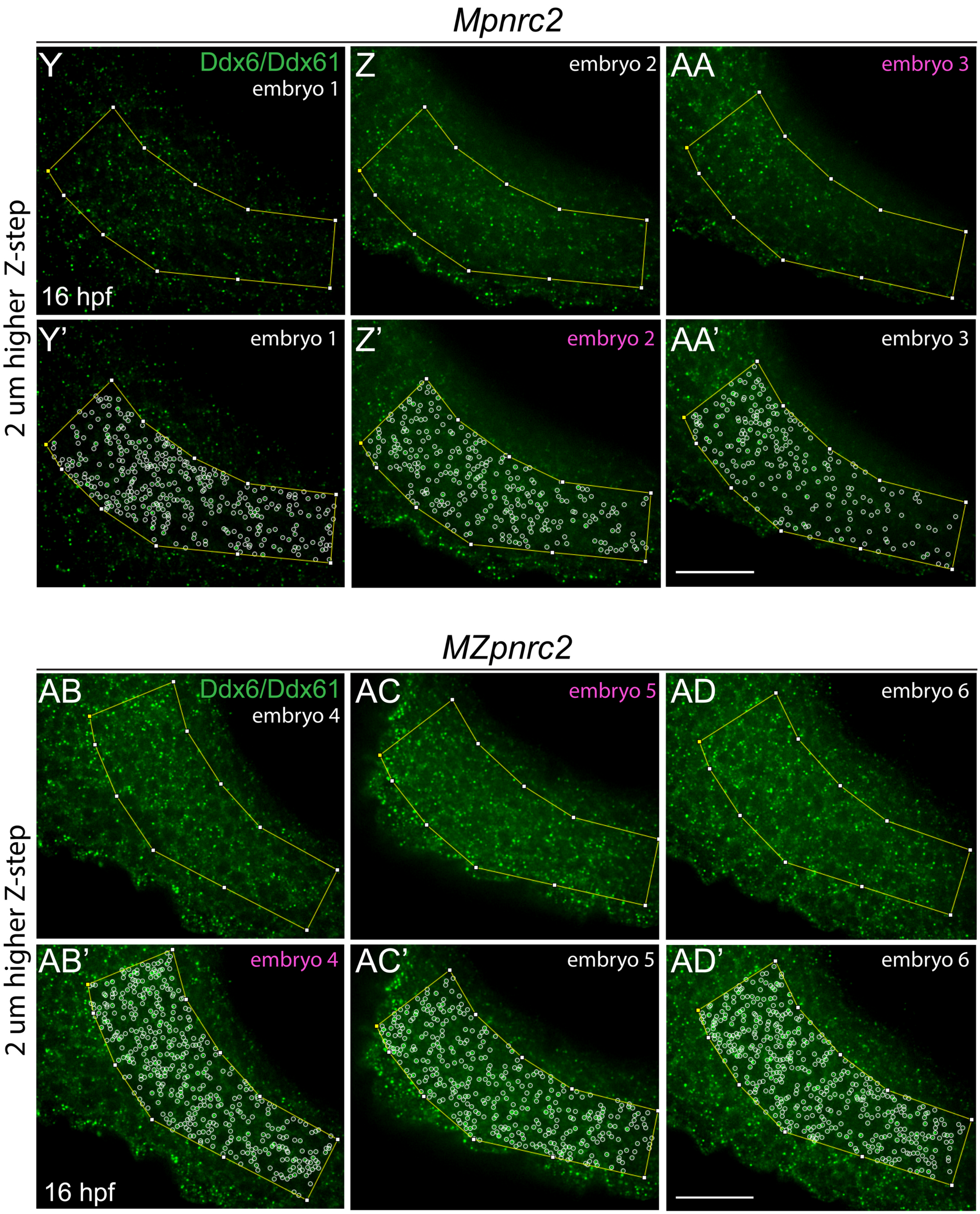

Supplementary Figure Legends

Fig. S4 (goes with Fig. 5). Ddx6/Ddx61 protein levels and Ddx6/Ddx61-containing puncta are increased in *MZpnrc2* mutant embryos. (A-L) Three z-stack slices, separated by 2 um intervals per slice, were analyzed for each embryo to comprehensively analyze Ddx6/Ddx61 immunolabeling in the presomitic mesoderm (PSM). Of the three slices, images and quantifications of lower and higher slices are shown (mean intensities in A-B and G-H; puncta numbers in D-E and J-K; respectively); quantifications of centrally-located z-stack slices (relative to higher and lower Z-steps) are shown in Fig. 5. Images with mean signal intensities closest to the mean for each genotype are shown in A-B and G-H with quantification in C and I, respectively. Images with puncta numbers closest to the mean of each genotype are shown in D-E and J-K with quantification in F and L, respectively. All quantifications were performed using three embryos per genotype. Error bars show standard deviation and p-values were calculated from an unpaired t-test. Embryo numbers (magenta text) indicate embryos with quantifications closest to the mean and correspond with magenta dots indicated in bar graphs (C,F, I, L). Images for all embryos analyzed are shown for lower Z-steps (M-R'), centrally-located Z-steps (S-X'), and higher Z-steps (Y-AD'). hpf = hours post-fertilization; scale bars = 50 um.

### Fig. S5 (goes with Fig. 7)

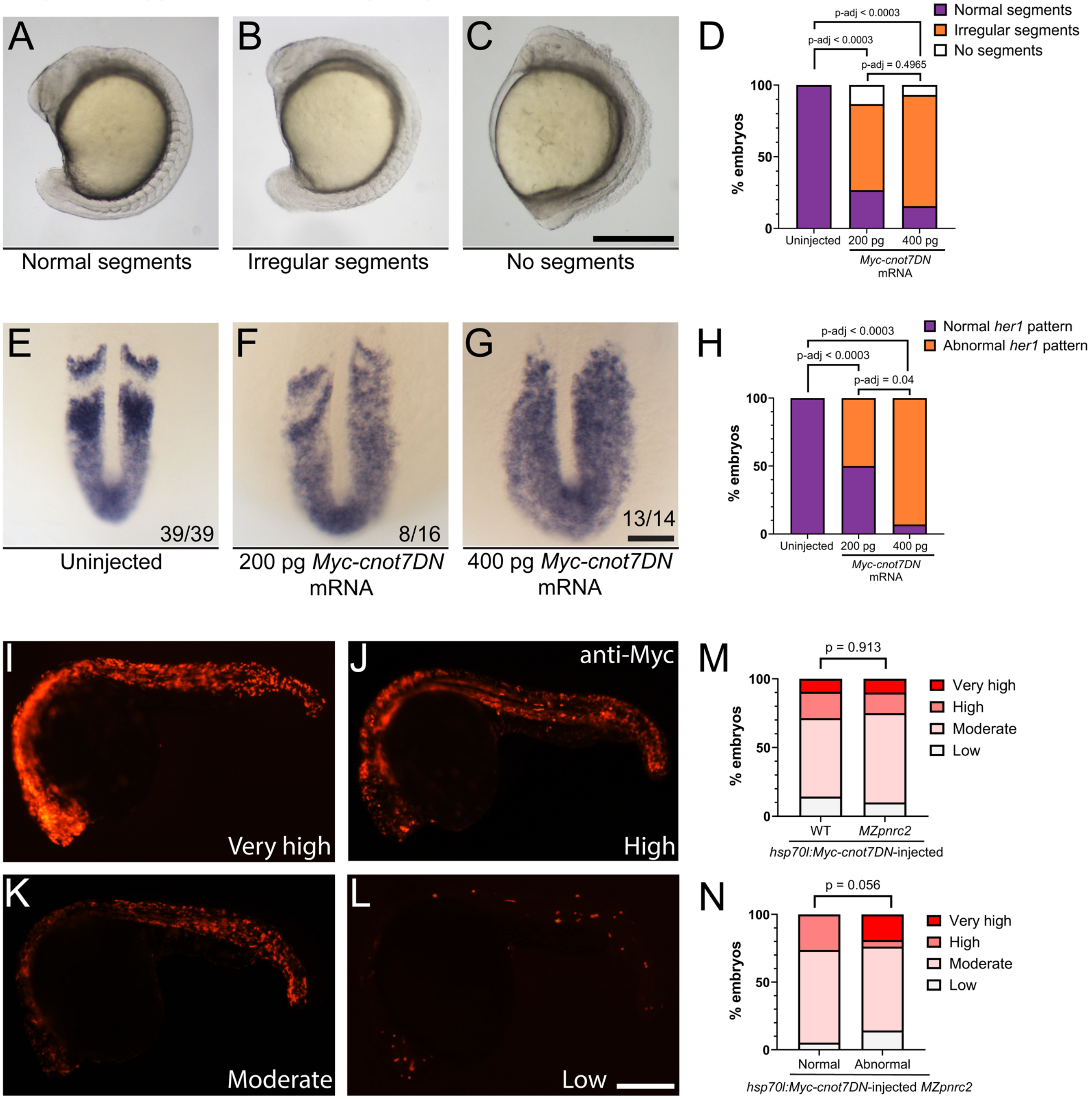

Fig. S5 (goes with Fig. 7). (A-H) Segmentation defects arise in a dose-dependent manner in WT embryos after injection of mRNA encoding a dominant negative deadenylase. WT embryos were injected with *Myc-cnot7DN* mRNA at the 1-cell stage and raised to mid-segmentation stages. Representative images are shown for embryos injected with *Myc-cnot7DN* mRNA with normal segments (A), irregular segments (B), and no segments (C). (D) Bar graph shows the proportion of embryos with normal, irregular, or no segments (n = 108 uninjected embryos; n = 30 embryos injected with 200 pg *Myc-cnot7DN* mRNA; n = 58 embryos injected with 400 pg *Myc-cnot7DN* mRNA). (E-G) A subset of uninjected and injected embryos shown in A-D were probed for *her1* expression by in situ hybridization. Representative embryos are shown for uninjected (E) and *Myc-cnot7DN* mRNA-injected embryos (F-G). (H) Bar graph shows the proportion of embryos with normal and abnormal *her1* expression patterns. Proportions were calculated from a single experiment and were consistent with a second independent experiment (not shown). (I-N) Induction of Myc-Cnot7DN expression after injection of a heat-shock inducible construct encoding a Myc-tagged dominant negative deadenylase. WT and *MZpnrc2* mutant embryos were scored for Myc protein expression after performing anti-Myc immunohistochemistry on plasmid-injected embryos (see methods in main text). Individuals were scored as having low (i.e. substantial mosaicism), moderate, high, or very high (i.e. little to no mosaicism) Myc expression on the basis of anti-Myc staining pattern. Representative embryos are shown for embryos in each expression class (I-L). (M) Bar graph shows the proportion of embryos in each expression class comparing WT (n = 33) and *MZpnrc2* mutants (n = 58), showing that there is no difference in Myc-Cnot7DN expression when comparing WT and *MZpnrc2* mutant embryos (p = 0.913). (N) Bar graph shows the proportion of embryos in each expression class comparing *MZpnrc2* mutants with normal (n = 25) and abnormal (n = 33) segments, showing that all *MZpnrc2* mutants with very high Myc-Cnot7DN expression (i.e. embryos with broad expression with little to no mosaicism) have abnormal segments, though overall, expression differences between *MZpnrc2* mutants with and without somite defects did not reach significance (p = 0.056). Proportions were calculated by combining data from two independent experiments that showed consistent results. P-values were calculated using a Fisher's Exact test comparing the number of embryos within each phenotype class. Adjusted p-values that correct for multiple pairwise comparisons were calculated using a Bonferroni adjustment (Bonferroni, 1935; Bonferroni, 1936; Dunn, 1961). hpf = hours post fertilization; DN = dominant negative; scale bars = 200 um (A-C & I-L), 50 um (E-G).

### Fig. S6 (goes with Figs 8-9)

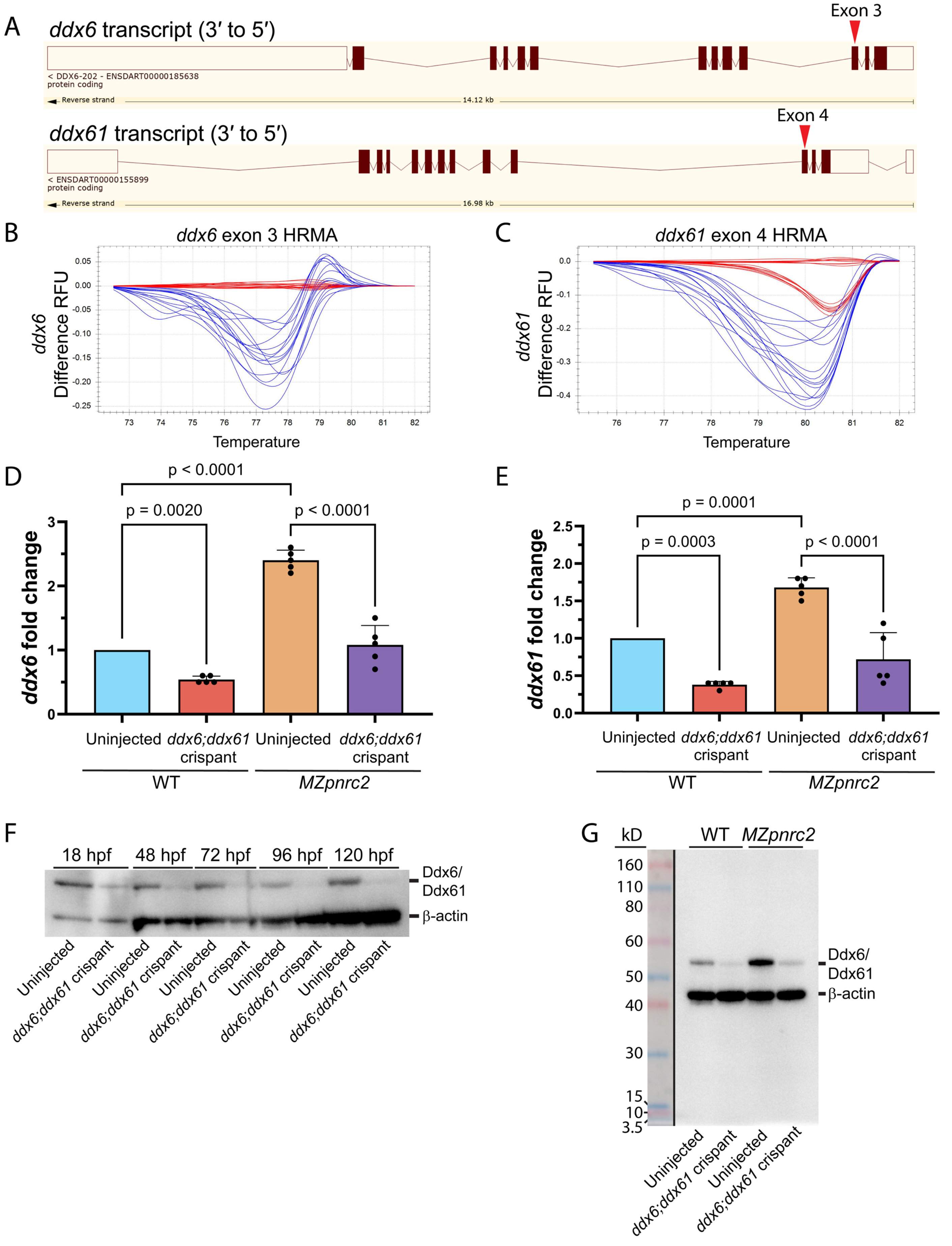

Supplementary Figure Legends

Fig. S6 (goes with Figs 8-9). CRISPRs targeting *ddx6* and *ddx61* efficiently reduce *ddx6* and *ddx61* mRNA and Ddx6/Ddx61 protein levels. (A) Cartoon schematics (exported from Ensembl) depict *ddx6* and *ddx61* gene structure with targeted regions denoted by red arrows, both of which lie just upstream of the DEAD-box helicase domain. (B-C) High resolution melt analysis (HRMA) difference plots showing CRISPR-induced mutagenesis at *ddx6* (B) and *ddx61* (C) targeted loci. CRISPR-injected embryo samples (blue) show strong deflections from uninjected embryos samples (red), indicating mutagenesis at the desired target loci. The presence of two distinct clusters (i.e. melt behavior patterns) in uninjected controls for *ddx61* (C) is due to a polymorphism outside of the targeted region within *ddx61* intron 3. (D-E) Bar graphs showing RT-qPCR results from 18 hpf embryos with fold change values derived from the mean of 5 biological replicates (biological replicate values derived from the average of 2 technical replicates each). Both *ddx6* and *ddx61* transcripts are significantly overexpressed in *MZpnrc2* embryos (D and E, blue vs orange bars), validating RNASeq results. Injection of *ddx6* and *ddx61* CRISPRs significantly decreases *ddx6* and *ddx61* transcript levels in both WT and *MZpnrc2* mutants at mid-segmentation (D and E, blue vs red bars and orange vs purple bars). Adjusted p-values calculated using a one-way ANOVA with a Dunn-Šídák correction for multiple comparisons (Šídák,1967; Abdi, 2007). (F) Immunoblot showing Ddx6/Ddx61 protein levels in WT uninjected and *ddx6*;*ddx61* crispant embryos over a 5 day period with β-actin as a loading control. (G) Full immunoblot image from Fig. 7G with molecular weight ladder indicating the expected sizes for Ddx6/Ddx61 and β-actin protein. Immunoblot results were consistent across two independent experiments. kD = kilodaltons.

Fig. S7 (goes with Fig. 9)

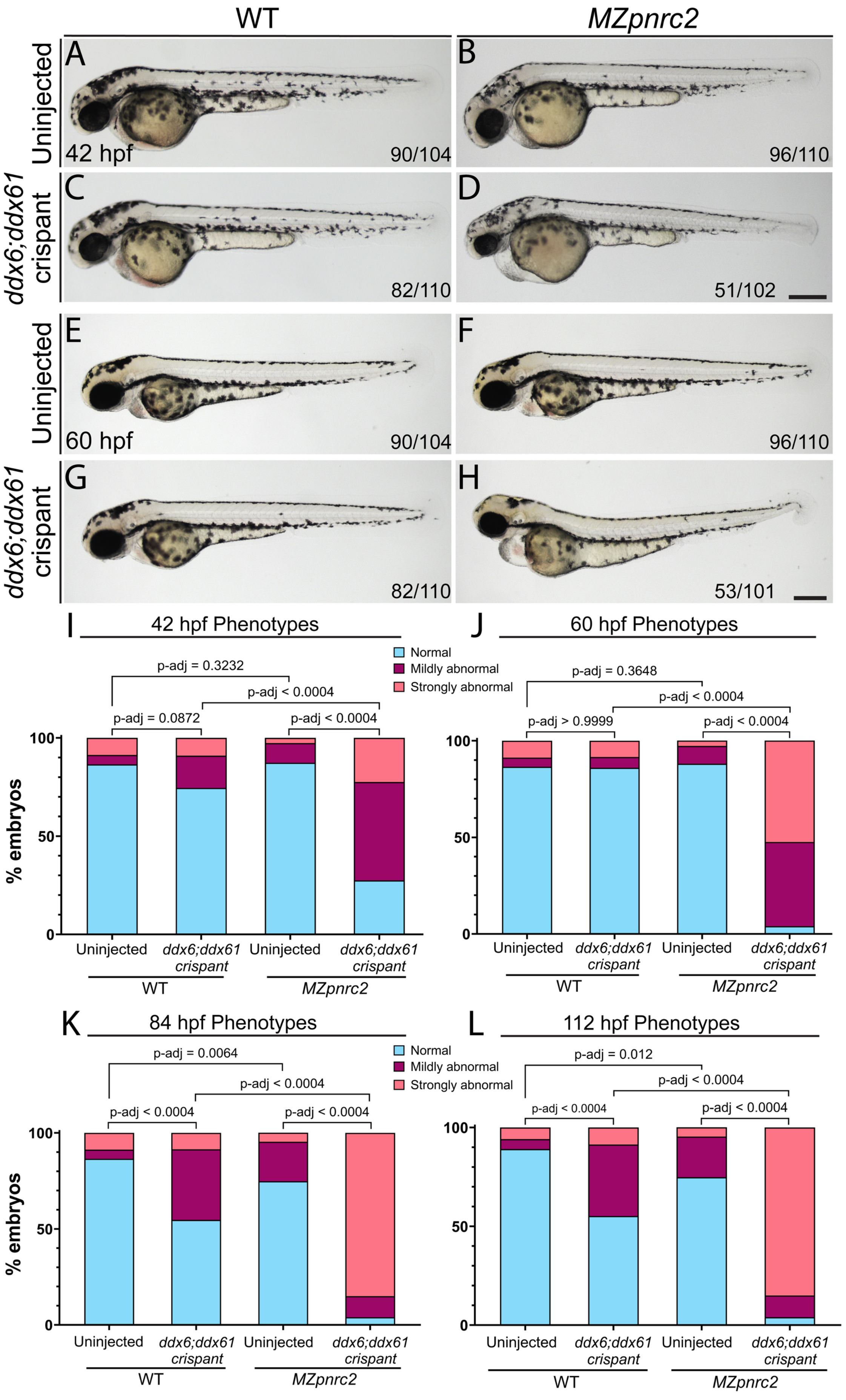

Supplementary Figure Legends

Fig. S7 (goes with Fig. 9). Abnormal phenotypes from depletion of Ddx6 and Ddx61 worsens over time in sensitized *MZpnrc2* mutants whereas WT embryos develop normally up to 60 hpf. (A-H) Representative embryos from the predominant class are shown at 42 hpf and 60 hpf, respectively, for (A, E) uninjected WT embryos and (B, F) uninjected *MZpnrc2* embryos, (C, G) *ddx6*;*ddx61* crispants (on the WT background), (D, H) *MZpnrc2 ddx6*:*ddx61* crispants. (I-L) Bar graphs show the proportion of embryos with normal and abnormal phenotypes at 42 hpf (I), 60 hpf (J), 84 hpf (K) and 112 hpf (L), indicating that *MZpnrc2 ddx6*;*ddx61* crispants are sensitized to Ddx6/Ddx61 depletion. Adjusted p-values were calculated using a Fisher's Exact test comparing the number of embryos within each phenotype class (normal, mildly abnormal, strongly abnormal) and corrected for multiple pairwise comparisons using a Bonferroni adjustment (Bonferroni, 1935; Bonferroni, 1936; Dunn, 1961). Representative larvae from predominant phenotypic classes at 84 hpf and 112 hpf are shown in Fig. 7A-E. hpf = hours post-fertilization; scale bars = 250 um.

**Supplemental Table Legends.** All supplemental tables are provided as downloadable Microsoft Excel files.

Table S1. RNA-Seq summary. Mapping results generated from Kallisto and STAR alignments of RNA-Seq sequencing reads from WT and *MZpnrc2* mutant embryos at mid-segmentation stages are shown in separate tabs (see methods for additional details).

Table S2. Kallisto normalized counts. Normalized read counts generated from Kallisto alignments of RNA-Seq sequencing reads from WT and *MZpnrc2* mutant embryos at mid-segmentation stages.

Table S3. Sleuth results and uORF analysis. Output data from differential expression analysis of WT and *MZpnrc2* mutant embryos using the program Sleuth (see methods for additional details). Overexpressed transcripts ($q < 0.05$), under-expressed transcripts ($q < 0.05$), and unaffected transcripts (log2FC -0.05 to 0.05) are indicated in separate tabs. uORF-containing transcripts that were significantly overexpressed ($q < 0.05$) are shown in a separate tab (see methods for criteria used to identify uORF-containing transcripts).

Table S4. Motif and GO Input lists. 3'UTR sequences of overexpressed ($q < 0.05$), under-expressed ($q < 0.05$), and unaffected transcripts (log2FC -0.05 to 0.05) used for motif analysis are listed in separate tabs (see methods for additional details). Sequences were obtained from the Ensembl GRCz11 zebrafish genome assembly using the tool BioMart. Gene IDs and UniProt IDs used for GO enrichment analysis are listed in a separate tab (see methods for additional details).

Table S5. Motif & GO enrichment results. Motif enrichment results of 3'UTR sequences from Table S4 using the program STREME are shown for overexpressed ($q < 0.05$), under-expressed ($q < 0.05$), and unaffected transcripts (log2FC -0.05 to 0.05) (see methods for details). GO enrichment results are shown in separate tabs for the GO aspects “biological process”, “molecular function”, and “cellular component” (see methods for details).

Table S6. Primer and gRNA sequences. Primer names and sequences that were used for RT-qPCR, poly(A)-tail length assays, in situ hybridization probes, CRISPR-induced mutagenesis, and cloning for the *Myc-cnot7DN* expression construct are listed in separate tabs (see methods for additional details).

#### Supplemental Methods

##### DNA extraction and high resolution melt analysis

Individual embryos were lysed in 50 ul 1X ThermoPol Buffer (NEB) at 95°C for 10 minutes, digested at 55°C for 1 hour using 50 ug Proteinase K (Thermo Fisher), followed by Proteinase K inactivation at 95°C for 10 minutes. CRISPR mutagenesis efficiency was analyzed by high resolution melt analysis (HRMA) on a CFX Duet instrument (Bio-Rad) using 1 ul of DNA extract as template in a 20 ul reaction with Precision Melt Mix following standard manufacturer procedures (Bio-Rad). gRNA and primer sequences are listed in Table S6. Melt analysis was performed using Precision Melt Analysis Software v1.2 (Bio-Rad).

##### RNA extraction and cDNA synthesis

Whole embryos at mid-segmentation stages (n = 15 embryos per biological replicate for each genotype in Figs S2-S3) were solubilized in Trizol for RNA extraction and purified following standard procedures (Thermo Fisher). All genotypes and conditions were performed in biological triplicate. 500 ng of total RNA was reverse transcribed using Superscript IV reverse transcriptase and random primers (Fig. S2) or oligo(dT) primers (Fig. S3) and according to the manufacturer's instructions (Thermo Fisher).

##### Quantitative RT-PCR analysis

Quantitative RT-PCR was performed using PowerUp SYBR Green Master Mix (Thermo Fisher) and 4.5 ul cDNA (diluted 1:50 in Fig. S2, diluted 1:25 in Fig. S3) in 20 ul reactions, following manufacturer's procedures. Negative controls lacking template were included for each primer set. All reactions were subjected to thermal melting to confirm that each reaction gave single peaks. Transcript levels were normalized to *mobk13* (*mob4*) (Hu et al., 2016; Gangras et al., 2020; Tietz et al., 2020). Cycle thresholds (Ct) were determined using Bio-Rad CFX manager software. The average Ct of 2-3 technical replicates per sample was used for calculations. Changes in mRNA expression were calculated by $\Delta\Delta Ct = \Delta Ct\ target - \Delta Ct\ control$. Relative changes in mRNA expression levels are represented graphically as fold change, where relative mRNA fold change = $2^{-\Delta\Delta Ct}$. All graphs and statistics were generated using Prism 10 (GraphPad).

##### mRNA injection

The *Myc-cnot7DN* coding sequence from plasmid *pMA-Myc-cnot7DN* (see main methods) was PCR amplified and cloned into expression vector pCS2+ (Rupp et al., 1994; Turner and

Weintraub, 1994) using standard digestion-based cloning with the enzymes EcoRI-HF and XbaI (NEB) to generate the plasmid *pCS2-Myc-cnot7DN*. For deadenylation inhibition experiments, *Myc-cnot7DN* mRNA was synthesized using the SP6 mMessage Machine Kit (Thermo Fisher), diluted in 0.2M KCl with 0.1% phenol red, and injected into 1-cell stage embryos (400 pg mRNA per embryo). Primer sequences are listed in Table S6.

**Microscopy and imaging**

Live embryos were mounted in 3% methylcellulose and imaged at 63x magnification using AxioVision Software (Zeiss) on a MZFLIII Fluorescence Stereo Microscope (Leica) with an AxioCam HRc digital camera (Zeiss). See methods in main text for quantitative confocal microscopy.

**Protein extraction and Immunoblot analysis**

See methods in main text.
